# Data integration across conditions improves turnover number estimates and metabolic predictions

**DOI:** 10.1101/2022.04.01.486742

**Authors:** Philipp Wendering, Marius Arend, Zahra Razaghi-Moghadamkashani, Zoran Nikoloski

**Author notes:** authors contributed equally (Philipp Wendering, Marius Arend).

## Abstract

Turnover numbers characterize a key property of enzymes, and their usage in constraint-based metabolic modeling is expected to increase prediction accuracy of diverse cellular phenotypes. *In vivo* turnover numbers can be obtained by ranking of estimates obtained by integrating reaction rate and enzyme abundance measurements from individual experiments; yet, their contribution to improving predictions of condition-specific cellular phenotypes remains elusive. Here we show that available *in vitro* and *in vivo* turnover numbers lead to poor prediction of condition-specific growth rates with protein-constrained models of *Escherichia coli* and *Saccharomyces cerevisiae*, particularly in the ultimate test scenario when protein abundances are integrated in the model. We demonstrate that *in vivo* estimation of turnover number by simultaneous consideration of heterogeneous physiological data leads to improved prediction of condition-specific growth rates. Moreover, the obtained estimates are more precise than the available *in vivo* turnover numbers. Therefore, our approach provides the means to decrease the bias of *in vivo* turnover numbers and paves the way towards cataloguing *in vivo* kcatomes of other organisms.

## Introduction

Genome-scale metabolic models (GEMs) together with advances in constrained-based modeling have led to improved understanding of how cellular resources are used to fulfill different cellular tasks (Goelzer *et al*, 2015; Lerman *et al*, 2012; O’Brien *et al*, 2013). Recent advances are largely propelled by the development of protein-constrained GEMs (pcGEMs) in which the catalytic capacities of individual enzymes are linked to the allocation of enzyme abundances (Chen & Nielsen, 2021a). Such models have led to more accurate predictions of maximum specific growth rates on different carbon sources (Adadi *et al*, 2012; Beg *et al*, 2007; Sánchez *et al*, 2017), flux distributions (Sánchez *et al*, 2017), and other complex phenotypes (Malina *et al*, 2021) in *Escherichia coli* and *Saccharomyces cerevisiae*. However, the development of pcGEMs critically depends on integration of organism-specific enzyme turnover numbers, *k_cαt_*, comprising the kcatome of an organism (Nilsson *et al*, 2017).

Measuring the kcatome of an organism based on *in vitro* characterization is limited due to impossibility to purify specific enzymes, lack of availability of substrates, and knowledge of required cofactors, such that their relevance for studies of *in vivo* phenotypes remains questionable (van Eunen & Bakker, 2014; Labhsetwar *et al*, 2017). Proxies for turnover numbers, referred to as maximal *in vivo* catalytic rates, can be obtained by combining constrained-based approaches for flux prediction with measurements of protein abundance under different growth conditions or genetic modifications (Davidi *et al*, 2016; Heckmann *et al*, 2020; Xu *et al*, 2021). The results from this approach, that entails ranking of condition-specific estimates that use individual data sets, have shown that the proxies for *in vivo* turnover numbers generally concur with *in vitro k_cat_* values in *E. coli* (Davidi *et al*, 2016). However, applications with data from *S. cerevisiae* (Chen & Nielsen, 2021b; Heckmann *et al*, 2020) and *A. thaliana* (Küken *et al*, 2020) indicated that these proxies for *in vivo* turnover numbers do not reflect *in vitro* measurements. Another approach to estimate the kcatome relies exclusively on machine and deep learning methods that use variety of features of enzymes (e.g. network-based, structure-based, and biochemical) (Zikmanis & Kampenusa, 2012; Preprint: Li et al, 2021; Heckmann *et al*, 2018), resulting in predictive models that can explain up to 70% of the variance in turnover numbers obtained *in vitro*.

The estimates of turnover numbers are integrated into metabolic models by different constraint-based approaches that have been grouped into coarse-grained (e.g. MOMENT (Adadi *et al*, 2012), sMOMENT (Bekiaris & Klamt, 2020), eMOMENT (Wendering & Nikoloski, 2022), and GECKO (Sánchez *et al*, 2017)) and fine-grained (e.g. resource balance analysis (Goelzer *et al*, 2015; Lerman *et al*, 2012) and ME-models (O’Brien *et al*, 2013)). Of these, GECKO (Sánchez *et al*, 2017) has been adopted in several recent studies due to the elegantly structured formulation of the protein constraints. In addition, GECKO allows for integration of protein contents and correction factors that account for mass fraction of enzymes (*f*) included in the model as well as average *in vivo* saturation (*σ*) of all enzymes, facilitating the development of condition-specific models.

While data-driven estimation of *in vivo* turnover numbers improves the coverage of *k_cat_* values in pcGEMs, the available estimates usually lead to over-constrained models when using the allocation of total protein mass, not considered in flux balance analysis (FBA). In addition, the bias in the estimates of *k_cat_* values can be determined in the ultimate test, in which they are used alongside constraints from enzyme abundances to predict growth--a scenario that has not yet been examined. Here, we propose PRESTO (for protein-abundance-based correction of turnover numbers), a scalable constraint-based approach to correct turnover numbers by matching predictions from pcGEMs with measurements of cellular phenotypes – simultaneously – over multiple conditions. As a constraint-based approach, PRESTO facilitates the investigation of variability of the proposed corrections. We show that predictions of growth by pcGEMs of *S. cerevisiae* with turnover numbers corrected by PRESTO are more accurate than those based on the models that include *k_cat_* values corrected based on a contending heuristic that relies on enzyme control coefficients (Preprint: Domenzain et al, 2021). We also demonstrate that same conclusions hold when enzyme abundances are integrated into the *E. coli* model using PRESTO. Therefore, PRESTO paves the way to broaden the applicability of pcGEMs for organisms with biotechnological applications and to arrive at genotype-specific estimates of the kcatome.

## Results and Discussion

### PRESTO – protein-abundance-based correction of turnover numbers

For a given data set of protein abundances over a set of conditions, the enzymes with turnover numbers in a pcGEM can be partitioned into three groups. For instance, a data set of protein abundances that was recently used to estimate *in vivo* turnover numbers in *S. cerevisiae* (Chen & Nielsen, 2021b) includes 45%, 41%, and 14% measured over all, at least one (but not all), and none of the 27 used conditions, respectively. Therefore, there is then a different data support for correcting the *k_cat_* values of these classes of proteins. PRESTO relies on solving a linear program that minimizes a weighted linear combination of the average relative error for predicted specific growth rates and the correction of the initial turnover numbers integrated in the pcGEM model (Fig. 1, Methods). It further employs K-fold cross validation (here, K = 3) with 10 repetitions while ensuring steady state and integrating protein constraints for proteins measured over all conditions (Fig 1, Methods). The training set of conditions is used to generate a single set of corrected *k_cat_* values, that provide estimates of *in vivo* turnover numbers, by using the respective protein abundances. The resulting corrected *k_cat_* values are in turn used to determine the relative error of predicted specific growth rate for each condition in the test set using flux balance analysis with the pcGEM, while only applying a constraint on total protein content and measured uptake rates. The relative error of predicted specific growth rate along with the sum of introduced corrections are lastly used to select the value for the tuning parameter *λ* in the objective function of PRESTO, as done in machine learning approaches that rely on regularization.

**Fig 1.**
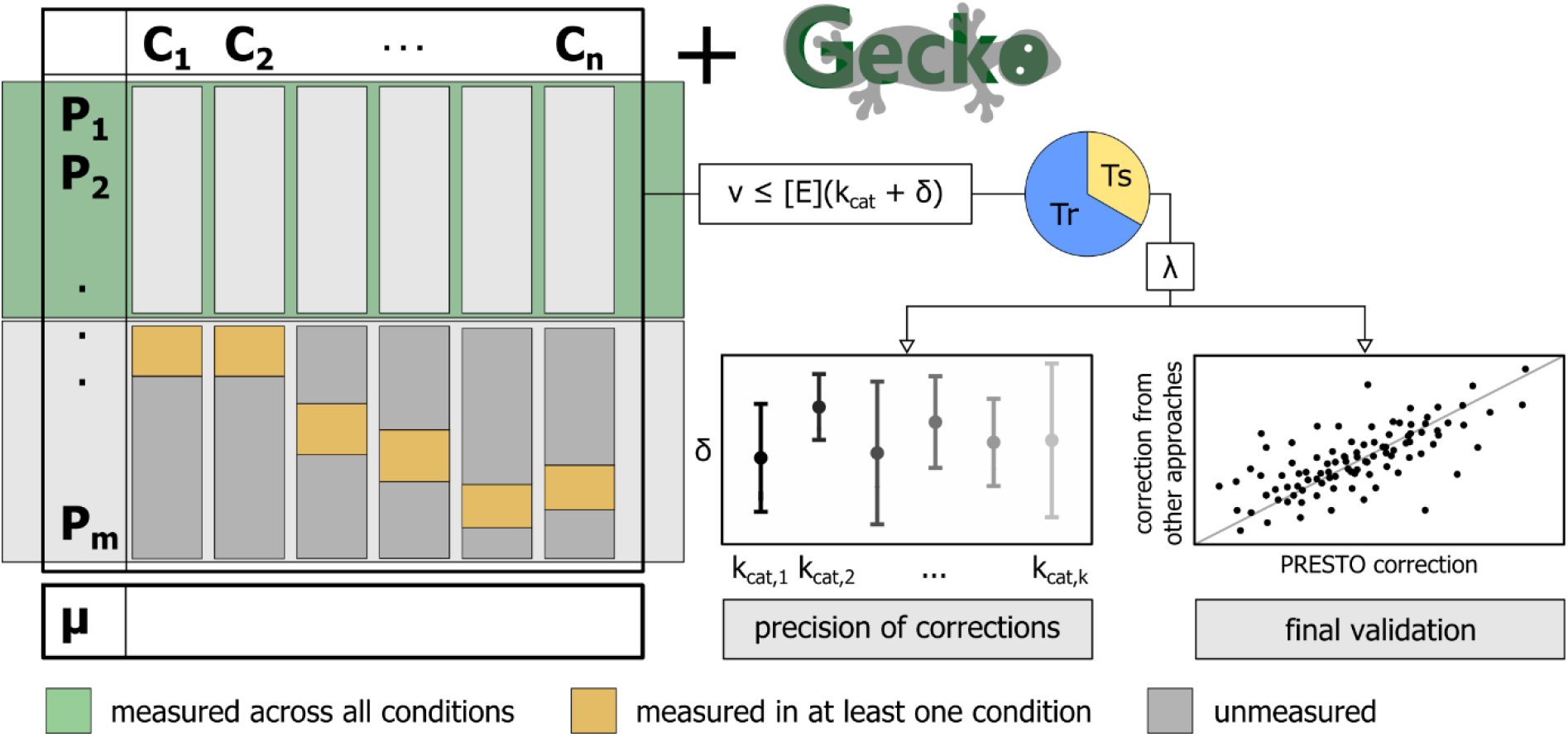
Schematic overview of the PRESTO approach for *k_cat_* correction. The approach uses a GECKO-formatted pcGEM (Sánchez *et al*, 2017) for an organism containing turnover numbers from BRENDA (Jeske *et al*, 2019). Using available data from *n* experimental conditions, *n* condition-specific models are generated using nutrient uptake rates and protein contents. PRESTO then uses data on abundances for the enzymes measured across the *n* investigated conditions and solves a linear program that minimizes a weighted sum of two objectives, the relative error to measured specific growth rates and the sum of positive *k_cat_* corrections, δ. The optimal weighting factor, λ, which modulates the trade-off between the two objectives, is then determined by cross-validation (Tr: training set; Ts: test set), choosing the parameter, which is associated with the lowest average relative error. Using the optimal value for λ, PRESTO combines all models for the experimental conditions to find a *k_cat_* correction for each enzyme with measured abundance. Last, the precision of δ values are assessed by variability analysis as well as by sampling and corrected *k_cat_* values are validated by comparing them to values obtained from other approaches.

### PRESTO outperforms a contending heuristic for correction of turnover numbers in *S. cerevisiae*

To determine the performance of PRESTO and compare it to that of contending heuristics, we used a data set comprising protein abundances and exchange fluxes from 27 diverse conditions, as supported by the principal component analysis (Fig S1). Application of PRESTO with a pcGEM of *S. cerevisiae* with initial *in vitro* turnover numbers obtained from BRENDA resulted in a mean relative error of 0.68 from the cross-validation procedure, yielding a correction of on average 213 turnover numbers (Fig S2a). For the *S. cerevisiae* pcGEM, we found a value of 10^−7^ for the parameter λ in the PRESTO objective provides the optimal trade-off between both the relative error and the sum of introduced corrections (see Methods). Moreover, we observed a high overlap between the sets of proteins with corrected turnover numbers in the cross-validation (average Jaccard distance of 0.07 (Fig S2b, c), suggesting that the integrated data from different conditions point at a specific subset of enzymes that need to be corrected to improve performance of growth prediction.

Unlike PRESTO, GECKO implements a heuristic for correction of turnover that is based on the control coefficient of a protein in each condition (Fig S3) (Preprint: Domenzain et al, 2021). The control coefficient of a protein is determined by increasing the turnover number by 1000-fold and scoring the effect on the predicted specific growth rate. The proteins are then ranked in a decreasing order of their control coefficients, and the turnover number of the first enzyme in the list is changed to the maximum value found in BRENDA for this enzyme across all organisms. This procedure is repeated with the remaining enzymes until the pcGEM predicts a growth rate that is at most 10% smaller than the measured specific growth rate for that condition or no additional *k_cat_* value that strongly constraints the solution can be found. In contrast to this procedure, PRESTO corrects at once the turnover numbers of multiple enzymes that are measured in all investigated conditions by simultaneously leveraging the data from the different conditions, considerably reducing runtime and the number of solved problems. As a result, rather than deriving condition-specific corrected *k_cat_* values, which are difficult to use in making predictions for unseen scenarios, PRESTO results in a single set of corrected *k_cat_* values.

Next, we compared the performance of PRESTO with the heuristic implemented in GECKO in three modeling scenarios that consider: (i) only condition-specific total protein content, (ii) both total protein content and uptake constraints, and (iii) additional constraints from abundances of enzymes measured in all conditions. For corrections of turnover number from PRESTO, we observed that the relative error spans the range from 0.15 to 0.88 in the least constrained scenario (i) (Fig 2a) and from 0.69 to 0.98 in the most constrained scenario (iii) (Fig 2c). In contrast, the relative error with the corrections of turnover numbers from the GECKO heuristic is in the range from 0.96 to 1.00 in scenario (iii) (Fig 2c). In addition, in scenario (iii), the relative error in the case of the GECKO heuristic for each condition is larger than the relative error of the PRESTO predicted specific growth rate (Fig 2c). We observed that predictions from FBA, considering enzyme abundances, without a constraint on the total protein content, led to an average relative error of 0.70 with *k_cat_* values corrected according to PRESTO and 0.99 with *k_cat_* values corrected according to GECKO (Table S1A).

**Fig 2.**
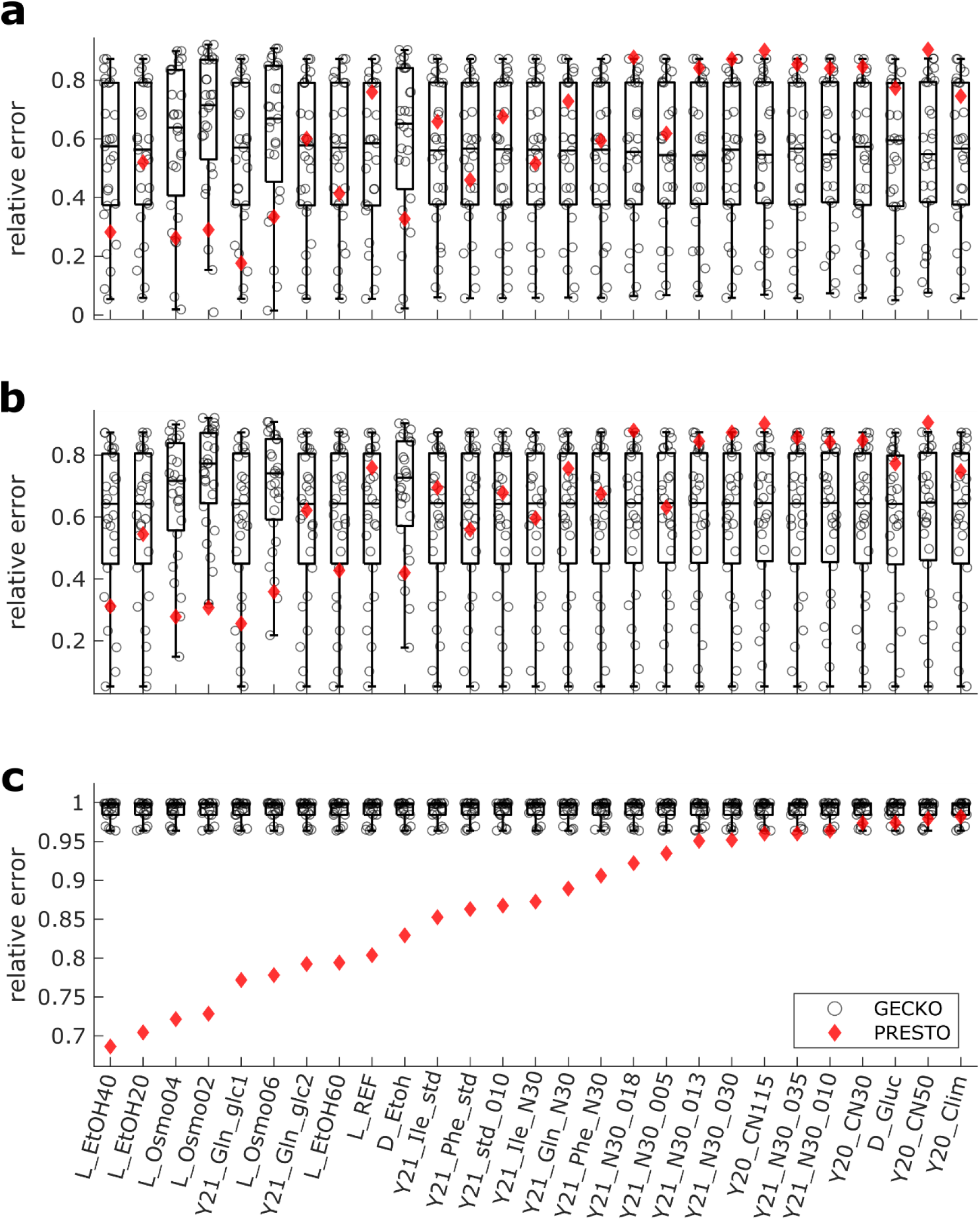
Comparison of predicted growth of *S. cerevisiae* from pcGEMs with *k_cat_* corrections from GECKO and PRESTO. Condition-specific pcGEMs with corrected *k_cat_* values generated in GECKO were used to predict the specific growth rate in each condition. The boxplots indicate the distribution of the relative error resulting from each set of condition-specific corrected *k_cat_* values obtained from the GECKO heuristic. Relative prediction error from each set is indicated by a circle. The red rhombus shows the relative error of predicted specific growth rate from the PRESTO model (*λ* = 10^−7^) by using the single set of corrected *k_cat_* values in the respective pcGEM. **(a)** Only the measured total protein pool was used to constrain the solution and condition-specific uptake rates were bounded by 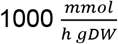; **(b)** measured uptake rates were also considered; **(c)** abundances of enzymes measured in all conditions were used as additional constraints. The compared pcGEMs in each condition used the same respective biomass coefficients, GAM, *σ*, and *Ptot* values (Methods). L: Lahtvee et al. 2017, D: Di Bartolomeo et al. 2020, Y20: Yu et al. 2020, Y21: Yu et al. 2021.

We also performed sensitivity analysis by investigating a smaller value, of 10^−10^, for the weighting factor λ used in the PRESTO objective. We found that when the weighting factor is 10^−10^ (at which the total corrections of the initial *k_cat_* values plateaus), the relative errors from PRESTO can be further decreased to 0.69 with the constraint on the total protein content, with no effects on the other findings (Fig S2a). We also note that the relative error lies in the range from 0.35 to 0.80 over the considered weighting factors in the range from 10^−14^ to 10^−1^. Together, these results demonstrated that *k_cat_* values corrected according to PRESTO provide better model performance than the values obtained by the contending heuristic in the case of *S. cerevisiae*.

### PRESTO provides precise corrections and estimates of *in vivo* turnover numbers

In the following, we investigated the precision of the corrected *k_cαt_* values from the application of PRESTO to data and pcGEM model of *S. cerevisiae*. To this end, we determined the range that the correction of the *k_cat_* value of each enzyme can take, while fixing the relative error in specific growth rate and total corrections from the optimum of PRESTO (Methods). Moreover, we complemented this analysis by sampling corrected *k_cat_* values that achieve the optimum of PRESTO with two values of the weighting factor λ of 10^−7^ and 10^−10^.

In the case of the corrected *k_cat_* values for *S. cerevisiae* with a weighting factor of 10^−7^, we found that the *k_cat_* values with the largest corrections are more precisely determined (Fig S4). In addition, the sampled corrections per enzyme show an average Euclidean distance to the respective mean of 4.88 s^−1^, indicating that the corrected values are more precise than the values in BRENDA, exhibiting an average Euclidean distance of 27.54 s^−1^ to the mean per EC number (Fig S5). Importantly, while *k_cat_* values with smaller correction showed larger variability, the 25 and 75 percentiles of the sampled corrections for 42 enzymes are concentrated around those resulting from PRESTO. Repeating the analysis with a weighting factor of 10^−10^ showed that the larger total corrections of the initial *k_cat_* values resulted in also larger variability for the corrections over all *k_cat_* (Fig S6). Here, too, for 62 enzymes the 25 and 75 percentiles of the sampled corrections are concentrated around those resulting from PRESTO. Therefore, we concluded that the corrections from PRESTO are precise and can be used in downstream analyses.

### Pathways of carbon metabolism are enriched in enzymes with corrected turnover numbers in pcGEM model of *S. cerevisiae*

In pcGEMs generated by the GECKO toolbox (Sánchez *et al*, 2017), turnover numbers are assigned to each enzymes in the GEM using a fuzzy matching algorithm. It takes into account the organism, substrate, and EC number of a BRENDA entry. When we investigated the magnitude of the turnover number correction dependent on the quality of the match between BRENDA entry and the corresponding enzyme, we found that *k_cat_* values measured in *S. cerevisae* were associated with smaller corrections then those from other organisms (Fig S7a).

To check which metabolic processes are more likely to require correction of *in vitro k_cat_* values, we next conducted an enrichment analysis based on the KEGG pathway terms linked to corrected *k_cat_* values (Methods). The most prominent pathway in this analysis was the synthesis of secondary metabolites, particularly the synthesis of cofactors and terpenoids (Fig 3a). However, several terms linked to central carbon metabolism, such as: the tricarboxylic acid cycle and oxidative phosphorylation, were also significantly enriched. Interestingly, amino acid synthesis was the only term linked to nitrogen metabolism that came up in this analysis, although many pathways of nitrogen metabolism were among the tested terms. This analysis suggested that particularly *in vitro* turnover numbers in carbon metabolism need to be corrected, due to underestimation in *in vitro* assays.

**Fig 3.**
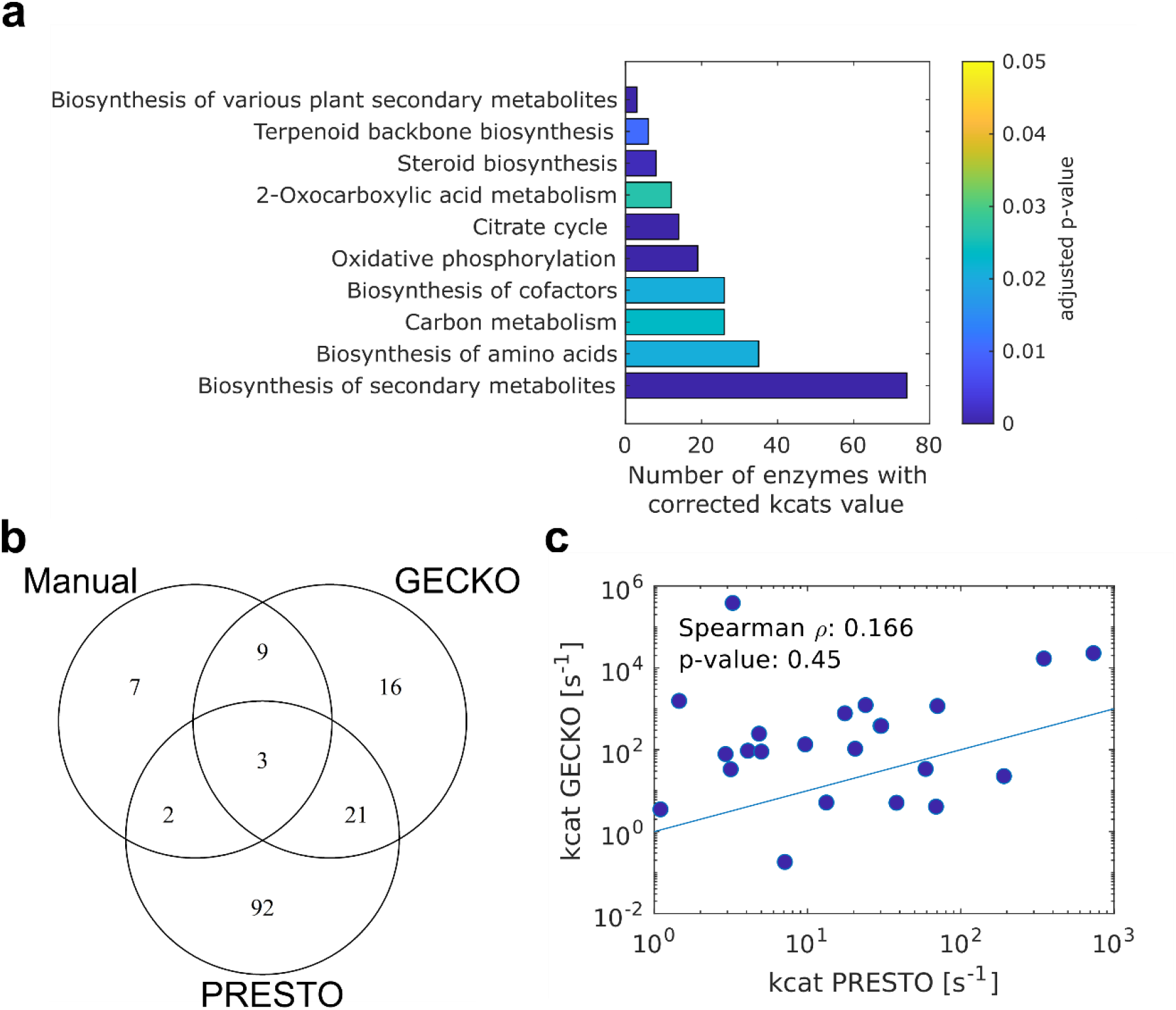
Comparison of enzymes with corrected *k_cat_* values by both GECKO and PRESTO. (a) KEGG Pathway terms significantly enriched in the set of enzymes corrected by PRESTO (*λ* = 10^−7^) in the *S. cerevisiae* model. The x-axis gives the number of corrected enzymes linked to the given term. The one-sided p-value was calculated using the hypergeometric density distribution and was corrected using the Benjamini-Hochberg procedure (Benjamini & Hochberg, 1995). **(b)** Venn diagram showing the overlap of enzymes whose *k_cat_* values were manually corrected (Sánchez *et al*, 2017) (“Manual”), automatically corrected by the GECKO heuristic in any of the conditions (“GECKO”), or corrected by PRESTO (“PRESTO”). **(c)** Log-transformed *k_cat_* values corrected using both the GECKO heuristic and PRESTO are not associated (Spearman correlation coefficient of 0.186, p-value = 0.385).

Next, we aimed to identify the extent to which the corrected *k_cat_* values differ between PRESTO and the GECKO approach. To this end, we determined the intersection of enzymes with *k_cat_* values corrected manually (Sánchez *et al*, 2017), by PRESTO, and by the GECKO heuristic. For this comparison, we considered the union of all condition-specific corrected *k_cat_* values from the GECKO approach. With the weighing factor *λ* = 10^−7^, PRESTO adapted the *k_cat_* values of 48% of enzymes corrected by the GECKO heuristic (Fig 3b, Table S1B). We did not find a significant Spearman correlation (*ρ_S_* = 0.17, *P* = 0.45) between the log-transformed *k_cat_* values in this intersection (Fig 3c), owing to the different principles employed in the two procedures. To determine the pathways that comprise enzymes whose turnover number are corrected by GECKO and PRESTO, we next repeated the pathway enrichment analysis for the enzymes in the overlap. Among the significant terms, like in PRESTO, we again found 2-Oxocarboxylic acid, amino acid, and secondary metabolism were enriched (Fig 3a, S8). However, the more specific pathway terms were associated with pathways that are part of carbohydrate metabolism and aromatic amino acid metabolism corrected by both approaches (Fig S8). In addition, the intersection between enzymes with manually corrected values and those corrected by the GECKO heuristic is higher than with PRESTO. This is expected since the manual curation partly aimed at correcting the most constraining turnover numbers (Sánchez *et al*, 2017).

We also compared the *k_cat_* values adjusted by GECKO against values for the maximum apparent catalytic rate obtained by parsimonious FBA (pFBA) using the same proteomics data (Chen & Nielsen, 2021b) (Fig S9 a, b). We confirmed the low correspondence (*ρ_S_* = 0.23) between the *k_cat_* values obtained from BRENDA, included in the GECKO model without manual modifications, and the maximum apparent catalytic rates. As expected, the correspondence of the maximum apparent catalytic rates to the turnover numbers corrected based on PRESTO was higher *ρ_S_* = 0.34). To investigate how turnover numbers obtained from pFBA perform as model parameters, we also generated a pcGEM in which BRENDA values were substituted by the maximum apparent catalytic rates from (Chen & Nielsen, 2021b), whenever available. In scenarios without enzyme abundance values, this model performed worse than that including the *k_cat_* values corrected by PRESTO as well as the model combining the maximum of all condition-specific GECKO corrections (Fig S10 a, b). In the enzyme abundance constrained scenario, the model with turnover numbers obtained from pFBA performed slightly better than GECKO but still only achieved a minimum relative error of 0.93, which is larger than 0.69 resulting from PRESTO (Fig S10c). These results demonstrated the value of the PRESTO in combining the genome-scale coverage of BRENDA with *in vivo* proteomics chemostat measurements to obtain less biased estimates of *in vivo k_cat_* values.

### Application of PRESTO with protein-constrained model of *E. coli* metabolism

To demonstrate the applicability of PRESTO across species, we deployed it with a pcGEM of *E. coli* (eciML1515) (Preprint: Domenzain et al, 2021; Monk *et al*, 2017). To this end, we used a large dataset comprising 31 different growth conditions (Valgepea *et al*, 2013; Peebo *et al*, 2015; Schmidt *et al*, 2016; Davidi *et al*, 2016). Due to the lack of data on nutrient exchange rates, the same GAM value (i.e., 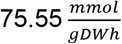) was used across all conditions. Similarly, we used the same value for total protein content since condition-specific measurements were not available (see Methods).

By applying a three-fold cross-validation, we found the optimal value for the λ parameter to be 10^−5^ (Fig S11a). This value was associated with an average relative error of 1.95 (overall average: 3.32) and 73 corrected turnover numbers, while on average 156 *k_cat_* values were corrected across all explored values for λ. On average, the Jaccard distance between cross-validation folds was 0.13 (Fig S11b), while the average Jaccard distance between unique sets of enzymes with corrected turnover numbers for each λ parameter was three-fold larger (0.4, Fig S11c). Thus, the corrected *k_cat_* values among cross-validation folds for each λ are more similar (maximum Jaccard distance of 0.29). Moreover, the union of the set of enzymes with corrected *k_cat_* values can remain similar over a range of chosen λ parameters up to four orders of magnitude (Fig S11c), demonstrating the robustness of the method.

The performance of PRESTO was assessed and compared to GECKO using scenarios (i) and (iii) since no condition-specific uptake rates were available. With default uptake rates, the relative error for predicted growth ranged between 0.01 and 8.56 in the less constrained scenario (i) (Fig 4a). Further, we obtained relative errors between 0.01 and 0.88 for the more constrained scenario (iii), when using the *k_cat_* values corrected by PRESTO (Fig 4b). In contrast, when using the *k_cat_* values from the GECKO approach, the relative error was in the range between 0.01 and the 4.89 for scenario (i) and between 0.89 and 0.99 for scenario (iii). In this scenario, too, we observed that the relative error using *k_cat_* values corrected by GECKO was consistently larger than the relative error resulting from the single set of corrected *k_cat_* values obtained by PRESTO (Fig 4a, b).

**Fig 4.**
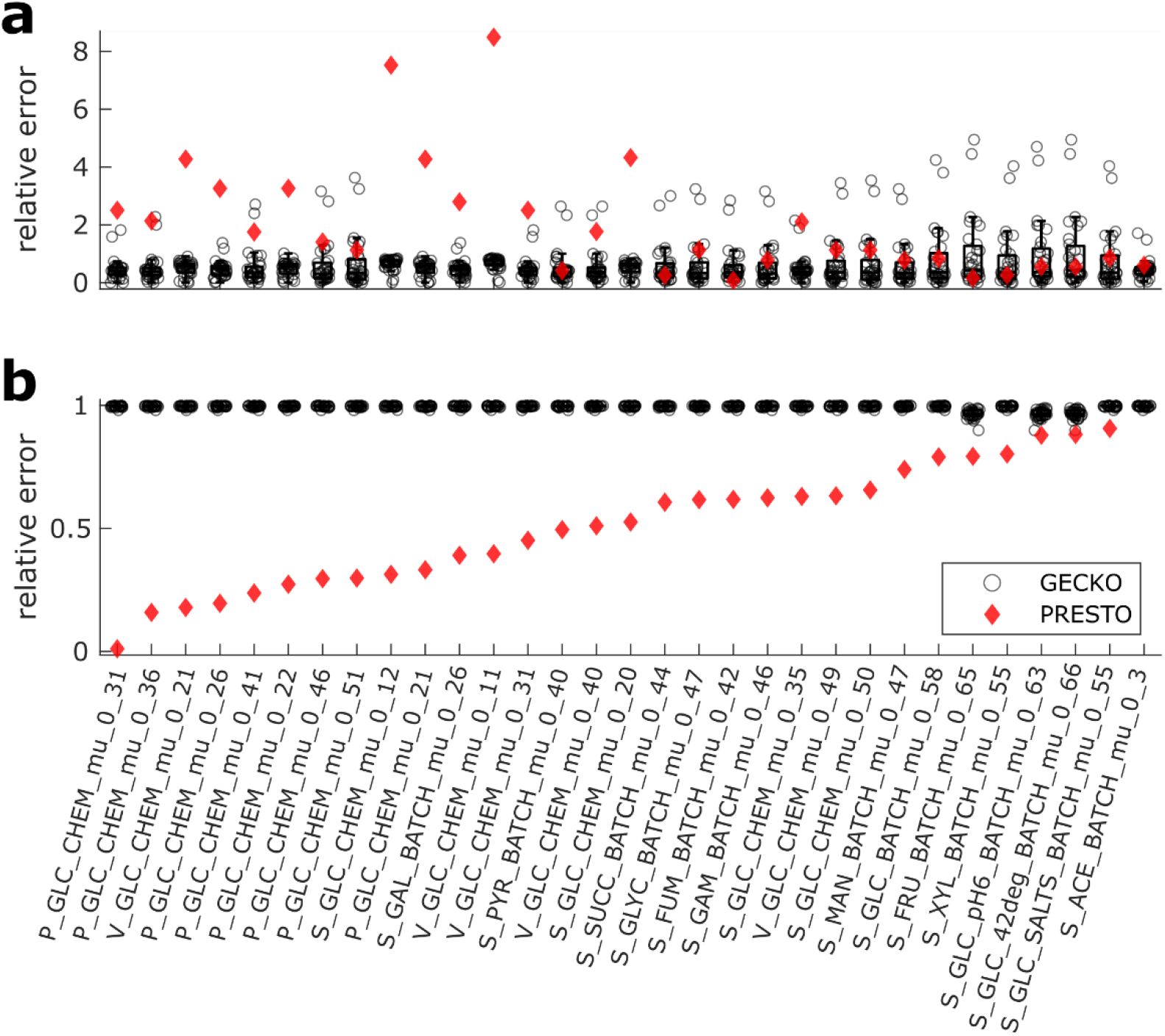
Comparison of predicted growth of *E. coli* from pcGEMs with *k_cat_* corrections from GECKO and PRESTO. Condition-specific pcGEMs with corrected *k_cat_* values generated in GECKO were used to predict the specific growth rate in each condition. Grey dots mark the relative error resulting from each set of condition-specific corrected *k_cat_* values obtained from the GECKO heuristic. Relative prediction error from each set is indicated by a circle. The red rhombus shows the relative error of predicted specific growth rate from the PRESTO model (Λ = 10^−5^) by using the single set of corrected *k_cat_* values in the respective pcGEM. **(a)** Only the measured total protein pool was used to constrain the solution and condition-specific uptake rates were bounded by 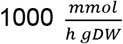; **(b)** abundances of enzymes measured in all conditions were used as additional constraints. Missing data points originate from infeasibility of the respective models. The compared pcGEMs in each condition used the same respective biomass coefficients, GAM, *σ*, and *P_tot_* values (Methods). P: Peebo et al. 2015, V: Valgepea et al. 2013, S: Schmidt et al. 2015.

The sum of introduced *k_cat_* corrections reached a plateau at 10^−11^ for the weighting factor *λ* in the PRESTO objective. We found that the relative cross-validation error at this value was 5.26, which is 2.7-fold larger than the relative error obtained using the optimal λ. Hence, allowing for more and larger corrections in PRESTO leads to a decrease of the overall relative error within the PRESTO program at the cost of highly biased parameters. The predictions with the highly biased parameters are worse in the test conditions and result in larger specific growth rate when no enzyme abundance constraints are enforced. This observation is in line with the small number of corrections introduced by the GECKO approach, where only the pool constraint is considered. We conclude that the prediction performance of the eciML1515 model was improved by using turnover numbers corrected by PRESTO only when enzyme abundances are integrated.

To assess the precision of the introduced *k_cat_* corrections, we performed variability analysis and sampling (see Methods) of the introduced corrections to the initial *k_cat_* values for two values of the weighting factor *λ*, namely 10^−5^ and 10^−11^. We observed that the 25 and 75 percentiles enclose a narrow interval around the values resulting from PRESTO (Fig S12) and are thus not evenly distributed across the respective interval determined by the variability analysis. We further noted that here, the predictions of smaller δ are generally more precise than the large corrections (*δ* ≥ *p*_50_), which span 2.12 orders of magnitude (small δ (< p_50_): 1.83, Fig S12). However, we also observed that the precision decreased when more corrections were allowed in PRESTO. This further justified our choice for the optimal parameter λ, which results in a lower number of 73 corrections compared to 204 at *λ* = 10^−11^, and moreover guarantees more precise estimates (Fig S13). In conclusion, the application of PRESTO is not limited to a single species but presents a versatile tool for the correction of turnover numbers across species.

In contrast to the observations made in *S. cerevisiae* we found that a model parameterized with the turnover numbers estimated by pFBA (Davidi *et al*, 2016) outperformed both PRESTO and GECKO in the modeling scenario where no enzyme abundance constraints are taken into account (Fig S10d). This is due to the fact, that pFBA, in contrast to PRESTO and GECKO, allows for the decrease of *in vitro k_cat_* values, in turn leading to more accurate predictions. However, in the scenario with enzyme abundance constraints PRESTO predicts specific growth rates closer to the experimental observation in 87% of the conditions (Fig S10e). Thus, in this scenario the integration of information from different modeling conditions achieved in PRESTO serves to obtain *in vivo k_cat_* value that perform better than the 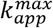 approach applied by (Davidi *et al*, 2016).

Interestingly, in contrast *S. cerevisiae*, we did not observe larger corrections by PRESTO for organism-unspecific *k_cat_* values (Fig S7b). The low number of corrections introduced by GECKO leads to an overlap of only three (75%) enzymes whose *k_cat_* values were also corrected by PRESTO (Table S1C, Fig S14a). These three enzymes catalyze reactions in three distinct metabolic pathways: Phosphoribosylformylglycinamidine synthase acts in the synthesis of purines, while serine acetyltransferase and NADP dependent Ketol-acid reductoisomerase are involved in the synthesis of sulfur aminoacids and hydrophic amino acids, respectively. The pathway enrichment analysis for all PRESTO corrections at *λ* = 10^−5^, indeed also identified amino acid and secondary metabolite synthesis as significantly enriched terms among the enzymes with corrected turnover numbers (Fig S14b). These results argue for a systematic underestimation of *in vivo* turnover numbers in these pathways compared to *in vitro* data, irrespective of the investigated organism. However, the lower order KEGG pathway terms enriched in *E. coli* do not overlap with the ones found in *S. cerevisiae*. Here, fatty acid metabolism and the synthesis of hydrophobic amino acids are among the pathways requiring correction of turnover numbers.

### Robustness of turnover number corrections

All of the approaches for estimation of turnover numbers rely on predicted (or estimated) fluxes and protein abundances from multiple conditions (Davidi *et al*, 2016; Chen & Nielsen, 2021b; Xu *et al*, 2021), but have not investigated the robustness of the estimates to the number of conditions used. Therefore, next, we investigate the difference in the sets of enzymes with corrected turnover numbers and the concordance of their corrections when ten randomly sampled subcollections of M experimental conditions (M = 3, 5, 10, 15) was used instead of all experiments. The differences and concordance were quantified with respect to the estimates obtained by considering data from all available experiments using the Jaccard index and the Pearson correlation coefficient, respectively. In the case of *S. cerevisiae*, we found that the smallest Jaccard difference over 200 scenarios was 0.36, while for *E. coli* this was 0.41 (Figs S15 and S16). In addition, the Pearson correlation coefficient between the (log-transformed) corrected turnover numbers with consideration of all versus a subcollection of M experiments in *S. cerevisiae* ranged from 0.99 to 1.00 (for M = 15) to 0.11 and 1.00 (for M = 3) (Fig S15). Repeating the analysis in the case of *E. coli*, we found that that the Pearson correlation coefficient ranged from 0.15 to 1.00 (for M = 15) to 0.14 and 1.00 (for M = 3) (Fig S16). This is in line with the expectation that the corrections stabilize with increasing number of experiments. Altogether, these findings pointed out the robustness of turnover number corrections derived from PRESTO with the number of available experiments.

## Conclusion

Characterization of enzyme parameters that can inform models of reaction rates is key to expanding and further propelling the usage of metabolic models in diverse biotechnological applications. While the generation of pcGEMs has facilitated the integration of more biophysically relevant constraints, it necessitates access to estimates of turnover numbers as key enzyme parameters. The bias in the available *in vitro* and *in vivo* turnover numbers can readily be assessed by considering the accuracy of growth predictions based on the integration of protein abundances. Indeed, we showed that condition-specific growth rates cannot be reliably predicted with pcGEMs of *S. cerevisiae* and *E. coli* when available *in vitro* and *in vivo* estimates of turnover numbers are used.

We use the modeling scenario that considers measured protein abundances as the ultimate test scenario not only for the prediction of metabolic fluxes, but also for the prediction of specific growth rates as it contains considerably more biochemically relevant constraints. The usage of protein abundances for estimation of proxies for *in vivo* turnover numbers also warrants their integration in pcGEMs to predict specific growth rates. However, we found that the integration of protein abundances resulted in poor predictions of specific growth rates. GECKO resolves this issue by flexibilizing measured protein abundances without considering physiological information during the procedure. To overcome this limitation, we developed PRESTO, which corrects turnover numbers and facilitates the integration of enzyme abundance constraints.

In contrast to PRESTO, GECKO uses measured total protein content from a single condition to achieve specific growth rates in the process of correcting the turnover numbers. As a result, the corrected turnover numbers vary between different experiments. Like in all existing approaches for estimation of turnover numbers based on GEMs, we integrated protein abundance data directly to correct turnover numbers. Following this strategy in PRESTO is further justified by the observation that the turnover numbers included in pcGEMs are often neither from the same enzyme (i.e., EC number), substrate, nor organism. While *in vivo* turnover number estimates can be adjusted by considering recently proposed Bayesian statistical learning {Formatting Citation}, this approach has not considered protein abundance information from proteomics measurements.

To resolve this issue, we proposed PRESTO, a constraint-based approach that simultaneously considers heterogeneous physiological read-outs and enzyme abundance measurements to correct *in vitro* turnover numbers. Through a series of comparative analyses, we demonstrated that the *in vivo* estimates of turnover numbers from PRESTO ultimately increase the prediction accuracy of condition-specific growth for the two organisms when enzyme abundance data are integrated in the corresponding pcGEMs. We also showed that the maximal *in vivo* catalytic rates, obtained by ranking of condition-specific estimates that use proteomics and fluxomics data, are more highly (but modestly) correlated to estimates from PRESTO than to *in vitro* turnover numbers. Owing to the constraint-based formulation of PRESTO, we also determined the precision of the *in vivo* estimates of turnover numbers. Previous studies have shown that even for the well-studied model organism *Sacharomyces cerevisiae*, only 52% of enzyme turnover numbers in the pcGEM can be obtained from organism-specific *in vitro* measurements (Preprint: Domenzain et al, 2021). Using organism unspecific *k_cat_* values for parameterization and correction of pcGEMs, as done in the GECKO pipeline, assumes that enzyme kinetic properties are comparable within one EC number class (Bar-Even *et al*, 2011; Davidi *et al*, 2018). However, we did not identify clear differences between EC classes, down to the second digit, when considering the distribution of *k_cat_* similarities within EC classes (Fig S17). Indeed, it has been reported that EC class plays only a minor role in the prediction of turnover numbers (Heckmann *et al*, 2018) and show stronger similarity with concordant GO categories (Mao & Ma, 2019). Interestingly, our findings show that the *in vivo* turnover numbers obtained from PRESTO are more centered around the means of the of the EC classes for both studied organisms (Fig S5). Together, these findings demonstrated PRESTO can be readily used to decrease the bias of *in vitro* and *in vivo* estimates of turnover numbers. This paves the way for employing the outcome of PRESTO and future extensions towards effectively predicting the kcatome from available protein sequences.

## Materials and Methods

### Experimental data

#### S. cerevisiae

We made use of a dataset gathered by (Chen & Nielsen, 2021b) from four different studies (Lahtvee *et al*, 2017; Yu *et al*, 2020, 2021; Di Bartolomeo *et al*, 2020), which included protein abundance data as well as measured growth or dilution rates and nutrient exchange fluxes. Exchange fluxes missing in certain conditions were set to 1000 mmol/gDW/h if the nutrient was present in the used culture media. We further augmented this data set by adding total protein content measurements from the original studies. For subsequent analyses, we used the maximum abundance of each protein over all replicates per experimental condition. Similarly, we used the average value for specific growth rates and nutrient exchange rates. Since no measurement of total protein content was available for the two conditions evaluated in the Di Bartolomeo study (Di Bartolomeo *et al*, 2020), we used the maximum protein content measured across the remaining conditions for these conditions (i.e., *0.67 g/gDW*). Moreover, we excluded three temperature stress conditions (i.e., Lahtvee2017_Temp33, Lahtvee2017_Temp36, Lahtvee2017_Temp38) from the analysis since temperature can have a large effect on the catalytic activity of an enzyme. Gene names in the proteomics dataset were translated to UniProt identifiers using the batch retrieval service of the UniProt REST API (Bateman, 2019).

#### E. coli

We used a dataset comprising 31 experimental conditions, which was gathered by Davidi and colleagues and augmented by Xu et al. (Davidi *et al*, 2016; Xu *et al*, 2021) from three publications (Valgepea *et al*, 2013; Schmidt *et al*, 2016; Peebo *et al*, 2015). Here, too, we used the maximum protein abundance over all replicates. Due to the absence of total protein content measurements in two of the original studies, we relied on the maximum protein content measured in the Valgepea study (i.e., 0.61 *g/gDW*) to be used for all conditions. Since precise data on nutrient uptake rates were only given for a few conditions, we assigned a default upper bound of 1000 mmol/gDW/h to all nutrients contained in the minimal medium (Table S2) with additional carbon sources as specified. Gene identifiers were translated to UniProt similar as for *S. cerevisiae*.

### Model preparation

The proposed approach aims at parsimonious correction of turnover values in genome-scale enzyme-constraint metabolic models using measured protein abundances. Therefore, it is important to consider differential association between enzymes and reactions, i.e., isozymes, enzyme complexes, and promiscuous enzymes. We decided to use the GECKO formalism (Sánchez *et al*, 2017), which deals with these problems elegantly by directly encoding the required information into the stoichiometric matrix. The genome-scale metabolic models for *S. cerevisiae* (YeastGEM v.8.5.0) and *E. coli* (iML1515) were obtained from the yeast-GEM and ecModels GitHub repository, respectively [(Preprint: Domenzain et al, 2021; Lu *et al*, 2019), https://github.com/SysBioChalmers; accessed on 22.08.2021]. For subsequent steps, functions of the COBRA v3.0 toolbox (Heirendt *et al*, 2019) and GECKO2.0 toolbox (Preprint: Domenzain et al, 2021) were employed, of which several functions were adapted for our purposes.

To arrive at raw enzyme-constraint models for both organisms, the GECKO2.0 model enhancement pipeline was adapted to allow the *k_cat_* correction procedure to be omitted. Moreover, any manual corrections of turnover numbers were excluded from model generation. In the process of adapting the raw pcGEM to the respective experimental conditions for both organisms, the GAM value per condition was fitted using the *scaleBioMass* function of GECKO2.0, based solely on the condition-specific nutrient exchange rates, and returning the minimum 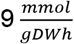) or maximum 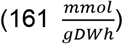 interval boundary if reached (only *S. cerevisiae)*. Furthermore, we omitted enzyme abundances, which were not measured across all experiments as the approach proposed here only works for enzymes with measured abundances (*E_measured_*).

### PRESTO approach

In the design of PRESTO, we modified the enzyme mass-balance constraints of the augmented stoichiometric matrix, created by GECKO, from

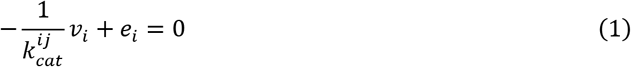

to inequality constraints that use the measured protein abundance directly and further assume a single turnover number per enzyme *i* over all catalyzed reactions 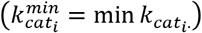:

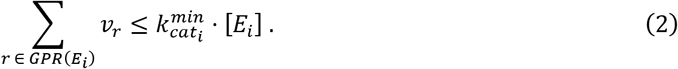

GPR stands for gene-protein-reaction rule that associates reactions with underlying genes and proteins. We justify making this assumption based on our observation that most enzymes in the *S. cerevisiae* model are associated with no more than four reactions. Further, the vast majority of enzymes are assigned a single unique turnover number even though they catalyze multiple reactions (Fig S16).

We then introduced a correction factor δ, which is added to each *k_cat_* if the protein abundances for the underlying enzyme were available:

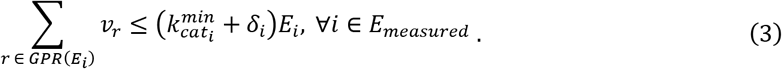

The value for each δ was constrained by the fold change *ε* with respect to 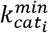 and a cut-off value *K^max^*, which denotes the maximum allowed *k_cat_* value.

To find a biologically relevant minimal set of adaptations with respect to the sum of δ, we minimized the weighted sum of the average relative error, ω, between measured (*μ^exp^*) and predicted specific growth rates (*ν_bio_*) over all experimental conditions *C*, and the average δ:

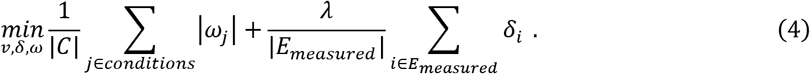

Finally, the linear programming formulation of the *k_cat_* correction in PRESTO is the following:

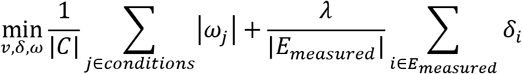

subject to

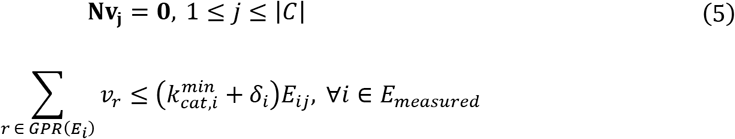

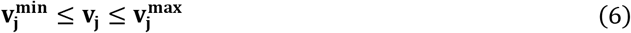

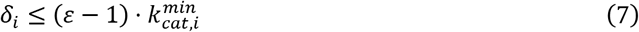

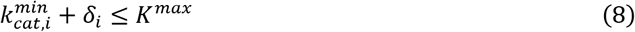

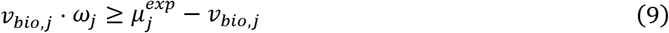

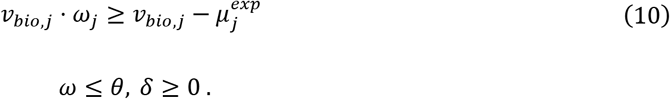

The value for ω was bound from above by a value θ, which was set to 0.6. Further, ε was set to 10^5^ since lower values did not yield solutions. The value for *K^max^* was set to 57,500,000 s^−1^(5.3.1.1, *Pyrococcus furiosus* (Sharma & Guptasarma, 2015)).

The parameter λ controls the trade-off between both minimization objectives (see Eq. 4). As λ is unknown and may also be condition- and model-specific, it was fitted using a 3-fold cross-validation scheme, which was repeated for 10 iterations. To this end, we scanned a log-scale interval between 10^−14^ and 10^−1^. In each iteration, we performed *k_cat_* corrections on two folds of condition-specific models and validated the obtained corrections on the remaining fold of condition-specific models. The relative errors (*e_r_*) and the sum of δ (i.e., Δ) were then used to calculate the scores *s_λ_*, which helped us choose the optimal value for λ:

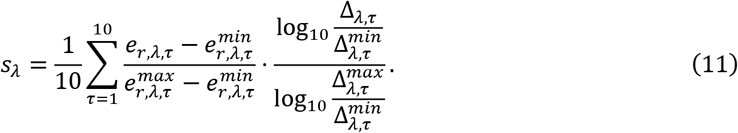

More precisely, the optimal value was then determined by finding the first sign change in the second numerical gradient over *s_λ_*, starting from the maximum value for λ. In addition to the optimal λ, we also compared our results to a second λ, where the sum of δ reached a plateau (*λ* = 10^−10^ for *S. cerevisiae* and *λ* = 10^−11^ for *E. coli*, Figs S2a and S10a).

### Variability analysis for δ

While PRESTO considers multiple experimental conditions to find a set of universal corrections for *k_cat_* values, it does not provide an exhaustive view over all possible solutions to this problem. To assess the precision of the corrections, we first performed a variability analysis for δ to find the minimum and maximum possible values. To guarantee that a solution of equal quality is found with respect to the previously determined sum of δ and the relative errors to experimentally measured specific growth rates (i.e., *ω^opt^*), corresponding constraints were added to arrive at the following linear programming problem:

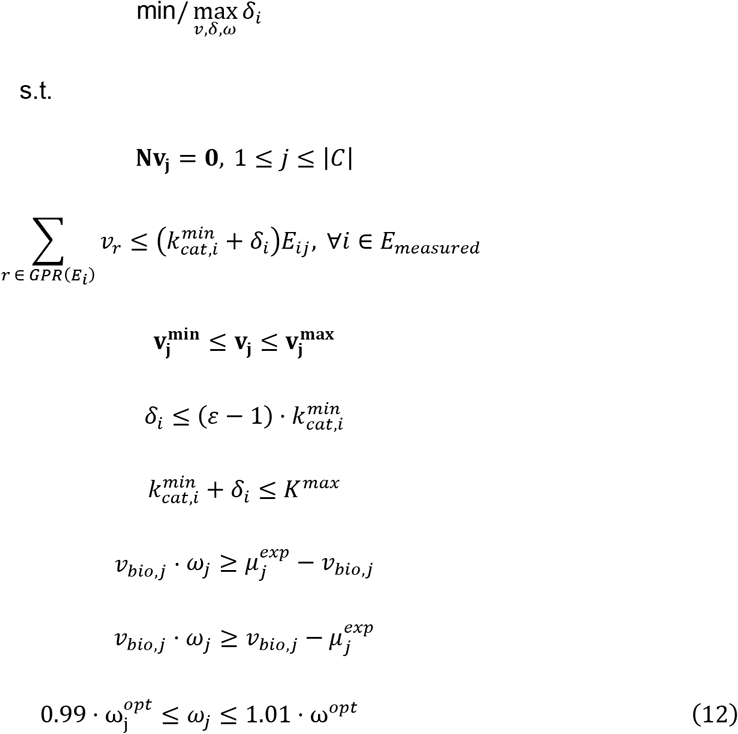

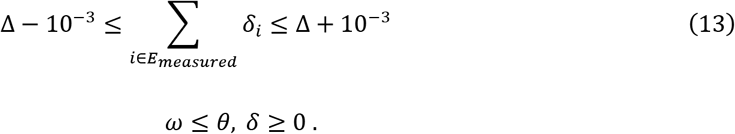

As the distribution within the obtained min/max intervals can be skewed, we sampled 10,000 points within the obtained intervals. For uniform random sampling, we created random vectors of corrections *δ*\* within the determined intervals and projected them onto the solution space by minimizing the distance of δ to the respective random vector. Therefore, we updated the objective of the program above:

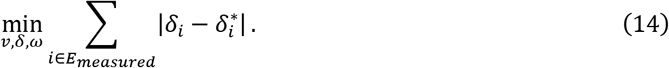

To ensure reproducibility and compatibility with the COBRA toolbox (Heirendt *et al*, 2019), we solved all optimization problems using the *optimizeCbModel* of the COBRA toolbox. Within this environment, we used the Gurobi solver v9.1.1 (Gurobi Optimization, 2021) but we note that any other supported solver can also be used. As we observed numerical instability of the problems in some cases, we decreased the feasibility tolerance (i.e., *feasTol* parameter) for the COBRA solver to 10^−9^ for all predictions.

### Validation of corrected models

We used the adapted GECKO pipeline (fitting a condition specific GAM; excluding manual *k_cat_* adaptions) to obtain models with *k_cat_* values adapted according to the objective control coefficient heuristic. We note that, when no manual modifications were introduced to the *Saccharomyces cerevisiae* models, the *k_cat_* adaption of the GECKO pipeline would stop because no objective control coefficient above the threshold of 0.001 could be found and corrected models would still be below the predicted growth error tolerance of 10%. To compare the predictive performance of PRESTO and GECKO corrected models, the models were adapted with the same condition-specific GAM, biomass reaction and total protein content, *P_tot_* Additionally, PRESTO models where constrained using the same condition-specific saturation rate, *σ* and enzyme mass fraction, *f*, as obtained from the GECKO pipeline. In contrast to the GECKO formulation, we did not subtract the mass of measured enzymes from the total protein pool constraint but instead introduced the measured protein concentration as upper bound on the enzyme usage reaction, *E_i_*, in the respective scenario. This formulation still guarantees that the mass of all used enzymes is lower or equal to the approximated cellular protein pool according to

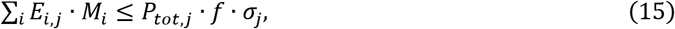

where *M* is the respective molecular weight of the protein. By considering measured and unmeasured enzymes in (15) we do not have to change *f* and use the same factor as for the scenario where no protein abundance measures are used (Sánchez *et al*, 2017). Maximum growth was predicted in three different constraint scenarios: (i) using only the protein pool constrain and default uptake rates (1000 mmol/gDW/h), (ii) using the pool constrain and experimentally measured uptake rates, (iii) using the previous constraints plus the absolute enzyme abundance.

The two studies which generated *in vivo k_cat_* estimates from pFBA (Davidi *et al*, 2016; Chen & Nielsen, 2021b)calculated a single value per reaction irrespective of the presence of isoenzymes. Thus, to parameterize the raw pcGEM (containing only uncorrected BRENDA values) we substituted the *k_cat_* values of all isoenzyme reactions with the same estimate provided in the study. Reactions catalyzed by complexes were not corrected. Since PRESTO and the pFBA studies provide a single condition independent model, we generated a condition independent GECKO model by following the maximum over all conditions approach: For the comparisons in Fig S9 the condition wise GECKO models were aggregated into a single union model in which for each reaction the maximum *k_cat_* value was used.

### Pathway enrichment analysis

The KEGG pathway terms (Kanehisa & Goto, 2000), associated with each enzyme that was measured in all conditions, were acquired using the KEGG REST API. The one-sided p-value, p, for significant enrichment of a pathway term among the enzymes with corrected *k_cat_* values was calculated using the hypergeometric density distribution:

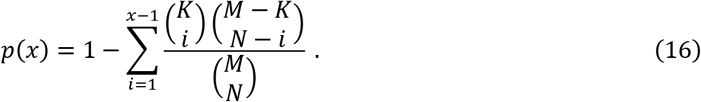

Only KEGG pathway terms associated to at least two corrected enzymes were taken into consideration. The p-values associated with all tested pathway terms were corrected for a false discovery rate of 0.05 using the Benjamini-Hochberg correction (Benjamini & Hochberg, 1995).

## Author Contributions

PW, MA performed research and analyzed data, PW contributed code for PRESTO approach, MA assessed model performance and performed statistical analysis, PW, MA, ZR, ZN designed research, PW, MA, ZN wrote the paper.

## Code Availability

All code that was used to generate the results of this study, including the PRESTO method, are available at https://github.com/pwendering/PRESTO.

