## Supplementary material for "Data integration across conditions improves turnover number estimates and metabolic predictions": SI Appendix

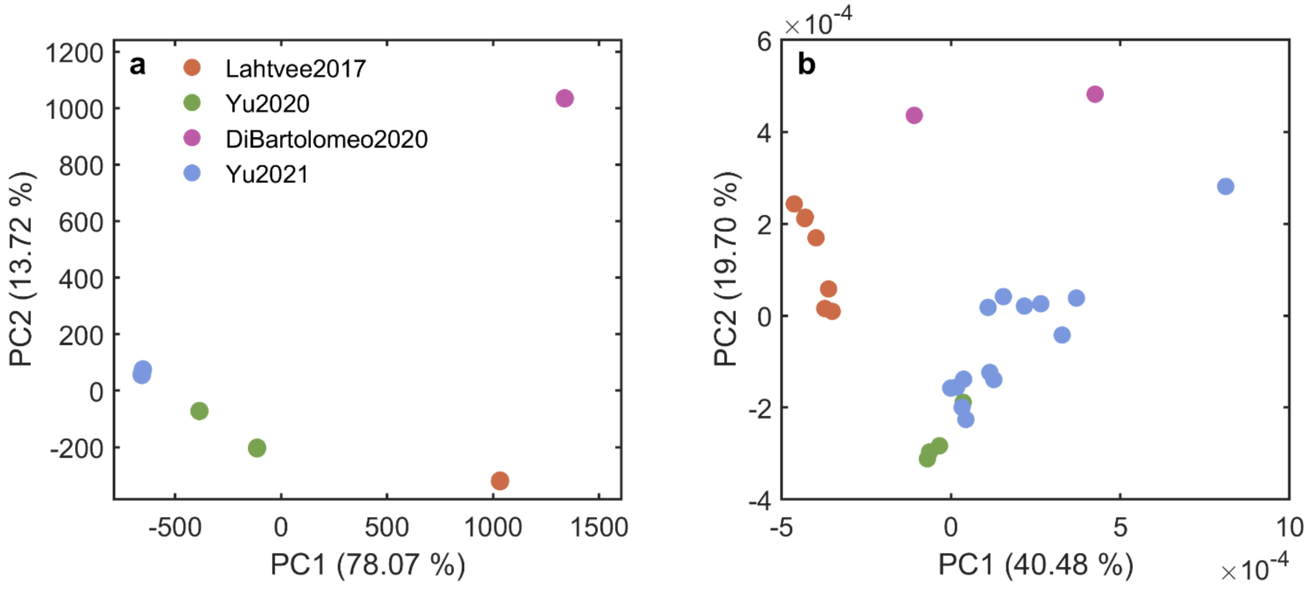


**Fig S1. PCA results for experimental conditions and protein abundances in the *S. cerevisiae* dataset.** Principal component analysis was performed using **(a)** nutrient exchange fluxes and total protein contents and **(b)** protein abundances from the 27 conditions used in the analysis. The colour of the points denotes the study from which the respective samples were obtained.

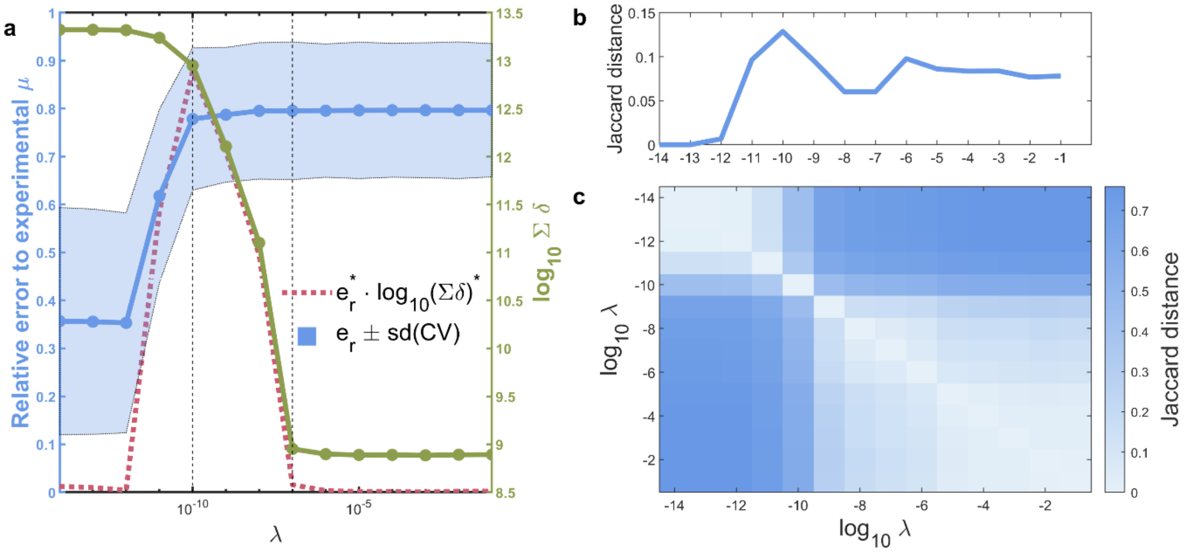


**Fig S2. Cross validation to obtain the optimal value for parameter λ using *S. cerevisiae* pcGEM**. **(a)** Shown are the average relative error ($e_r$, blue solid line), average sum of added corrections δ (green solid line), and the scoring metric that was used to find the optimal value (red dotted line). Altogether, 14 values for λ were explored in the range from $10^{-14}$ to $10^{-1}$. The optimal value for λ was determined by the first inflection point of the scoring metric (right dashed vertical line). The left dashed vertical line represents the value at which the sum of δ reaches a plateau. **(b)** Average Jaccard distance between cross-validation folds over ten iterations for each of the explored λ. **(c)** Average Jaccard distance between the union of corrected $k_{cat}$ values over ten iterations for each pair of explored λ values.

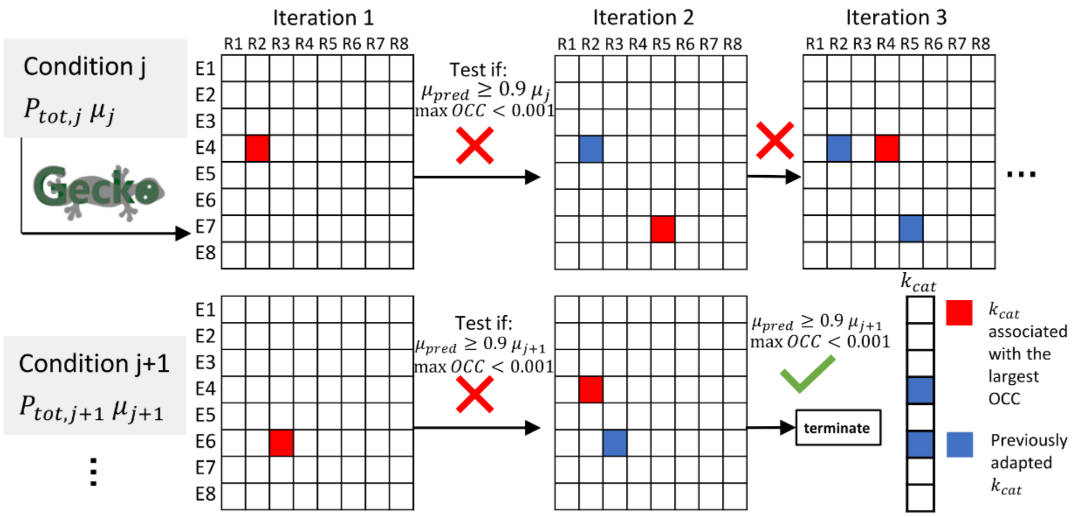


**Fig. S3. Schematic overview of the $k_{cat}$ correction process used in GECKO.** The tables correspond to simplified representations of the lower left part of the augmented stochiometric matrix produced by GECKO. To obtain condition-specific models, raw pcGEM models containing the turnover numbers matched from the BRENDA database are constrained using the condition-specific total protein content ($P_{tot}$). If the predicted growth rate from FBA ($\mu_{pred}$) is less than 0.9 of the experimentally observed growth rate ($\mu$) , each $k_{cat}$ value in the augmented stochiometric matrix is increased by 1000 independently and the effect on the growth rate is measured via the objective control coefficient (OCC). The $k_{cat}$ value associated with the highest OCC is set to the largest value associated with the respective EC number in the BRENDA data base. This procedure is iterated until the predicted growth rate is above the 0.9-fold threshold or no $k_{cat}$ values are associated with an OCC >0.001 and produces a set of condition specific $k_{cat}$ adaptions.

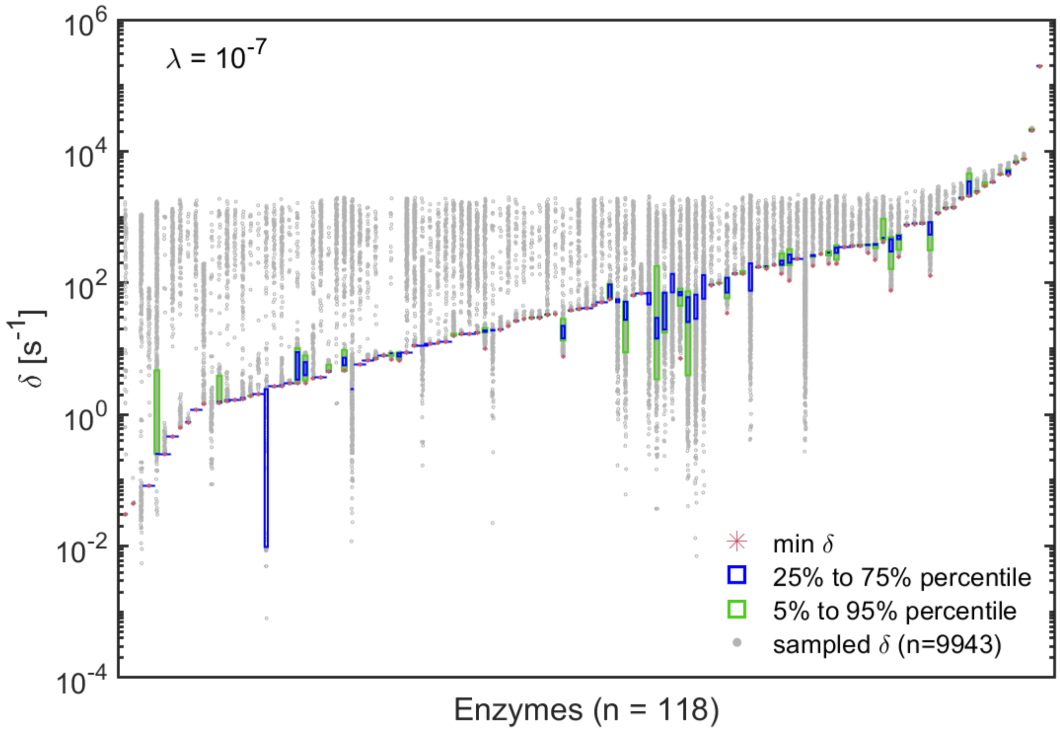


**Fig S4. Precision of $k_{cat}$ corrections introduced by PRESTO using the optimal λ value for the *S. cerevisiae* pcGEM.** The minimum and maximum values for each δ were determined by variability analysis in which the relative errors and the sum of corrections, δ, are fixed to values obtained from PRESTO. Using these ranges, 10,000 random points were sampled uniformly across the feasible region (9979 feasible linear programs).

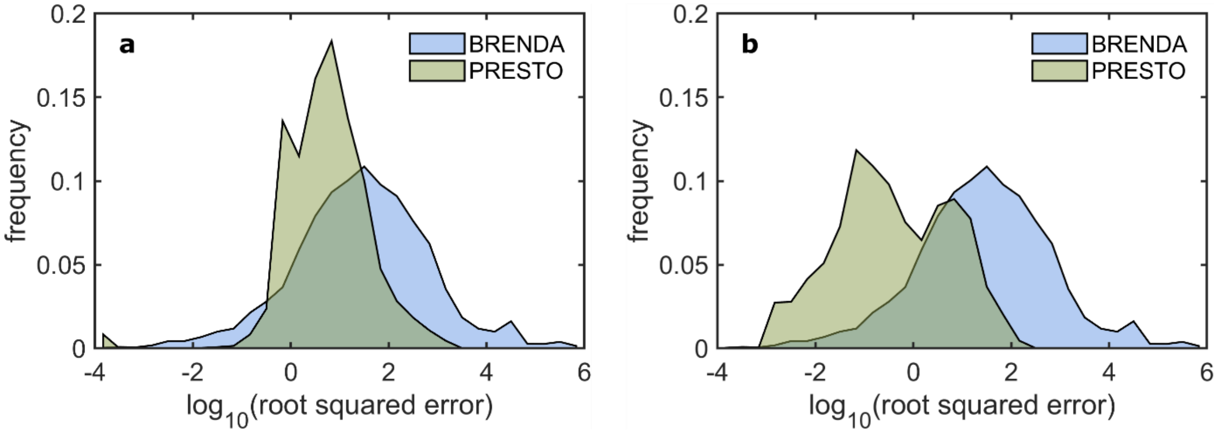


**Fig S5: Euclidean distance of corrections and $k_{cat}$ values from the respective mean per EC number / protein.** The root of squared distances to the average value of sampled corrections from PRESTO (per protein) and $k_{cat}$ values from BRENDA (per EC number). **(a)** shows the distributions for *S. cerevisiae* ($\lambda = 10^{-7}$) and **(b)** for *E coli* ($\lambda = 10^{-5}$). To allow for a fair comparison, we only used BRENDA EC numbers for which there are at least three entries for turnover numbers.

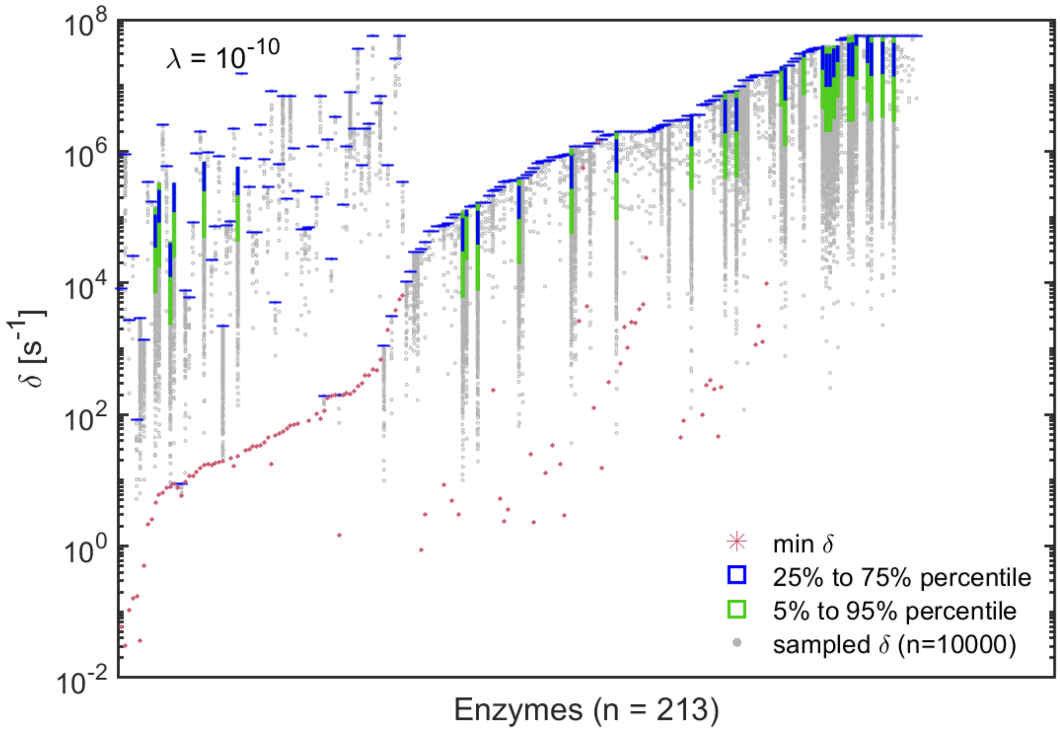


**Fig S6. Precision of $k_{cat}$ corrections introduced by PRESTO using the λ value for which the average sum of corrections plateaus in the *S. cerevisiae* pcGEM.** The minimum and maximum values for each δ were determined by variability analysis in which the relative errors and the sum of corrections, δ, are fixed to values obtained from PRESTO. Using these ranges, 10,000 random points were sampled uniformly across the feasible region. To arrive at feasible optimization programs, the feasibility tolerance for the solver was decreased to $10^{-6}$).

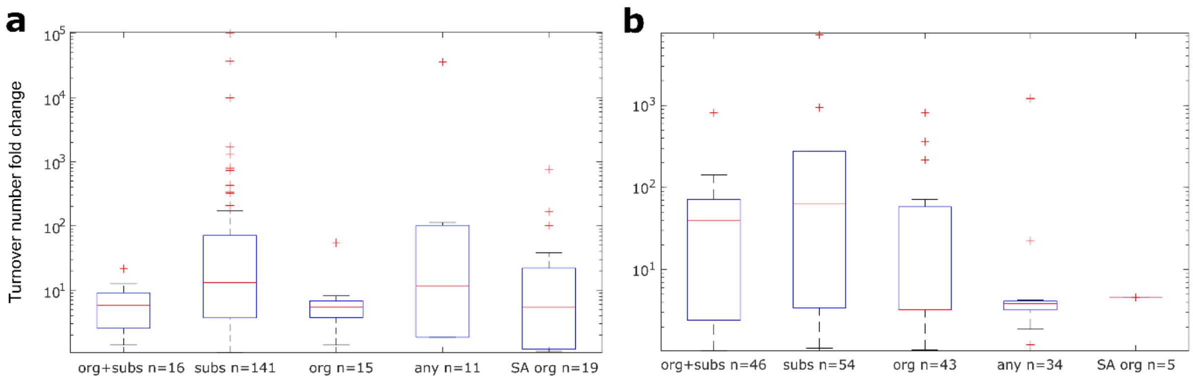


**Fig S7. Fold change of PRESTO turnover number correction for different GECKO $k_{cat}$ match qualities.** For reactions bearing turnover numbers corrected by PRESTO in (**a**) *S. cerevisiae* ($\lambda = 10^{-5}$) or **(b)** *E. coli* ($\lambda = 10^{-5}$), the quality of the initial BRENDA fit in the uncorrected GECKO model was extracted. The boxplots indicate the distribution of turnover number fold changes in the different matching categories. Categories indicated on the x-axis are exact match of organism and substrate (org+subs), exact match of substrate (subs), exact match of organism (org), only matching EC number (any), exact match of organism in the specific activity database (SA org). The group of $k_{cat}$ values matched only by EC number in the specific activity database did not have more than 4 entries in any of the organisms and was therefore omitted in this plot.

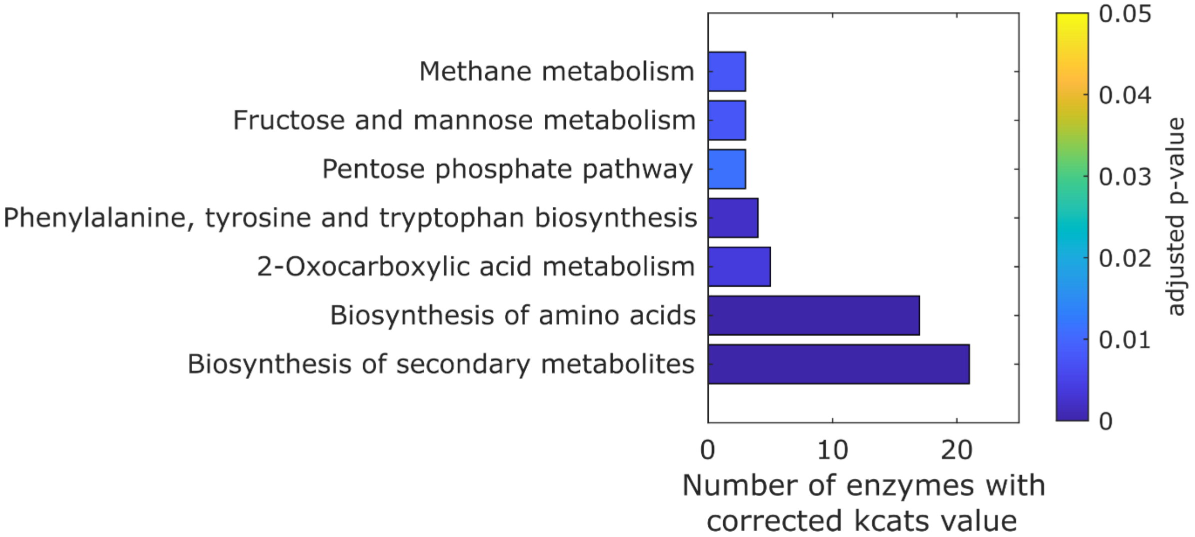


**Fig S8. KEGG pathway terms significantly enriched in the set of enzymes whose turnover numbers were corrected by GECKO and PRESTO in *S. cerevisiae.*** The x-axis gives the number of corrected enzymes linked to the given term. The total number of enzymes in the tested set was 24. Detailed info on the set can be found in Table S1D. The one-sided p-value was calculated using the hypergeometric density distribution and was corrected using the Benjamini-Hochberg procedure (Benjamini & Hochberg, 1995).

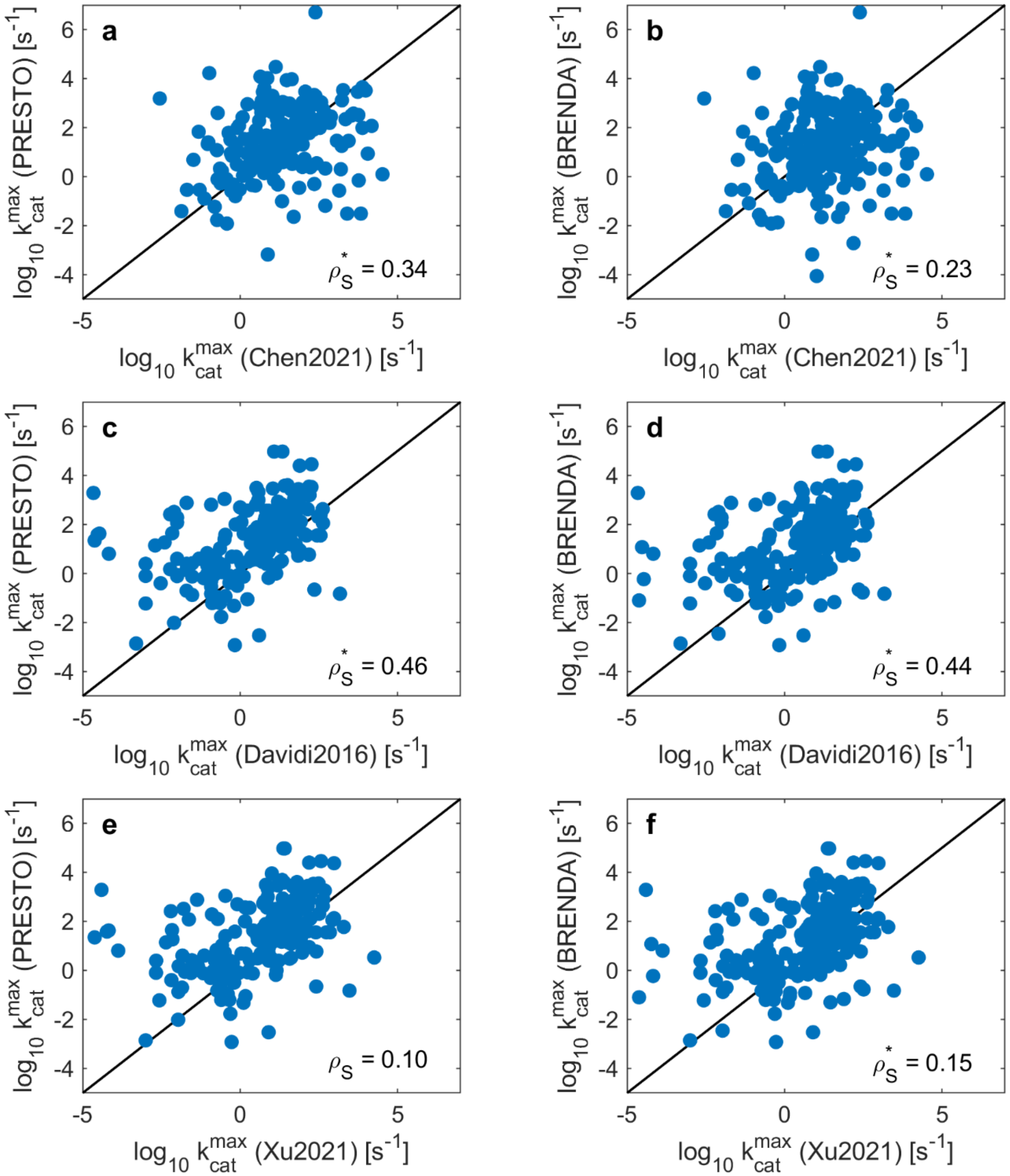


**Fig S9. Comparison of corrected $k_{cat}$ values from PRESTO with previously published $k_{app}$ values for *S. cerevisiae* and *E. coli*.** Comparison of maximum $k_{cat}$ values per reaction with predicted $k_{app}$ values from Chen et al. (2021) (*S. cerevisiae*; **a,b**),. Davidi et al. (2016) (*E. coli*; **c,d**), and Xu et al. (2021) (*E. coli*; **e,f**). The comparison was only performed for reactions associated to homomeric enzymes. The solid line represents hypothetical full correlation. $\rho_S$: Spearman correlation.

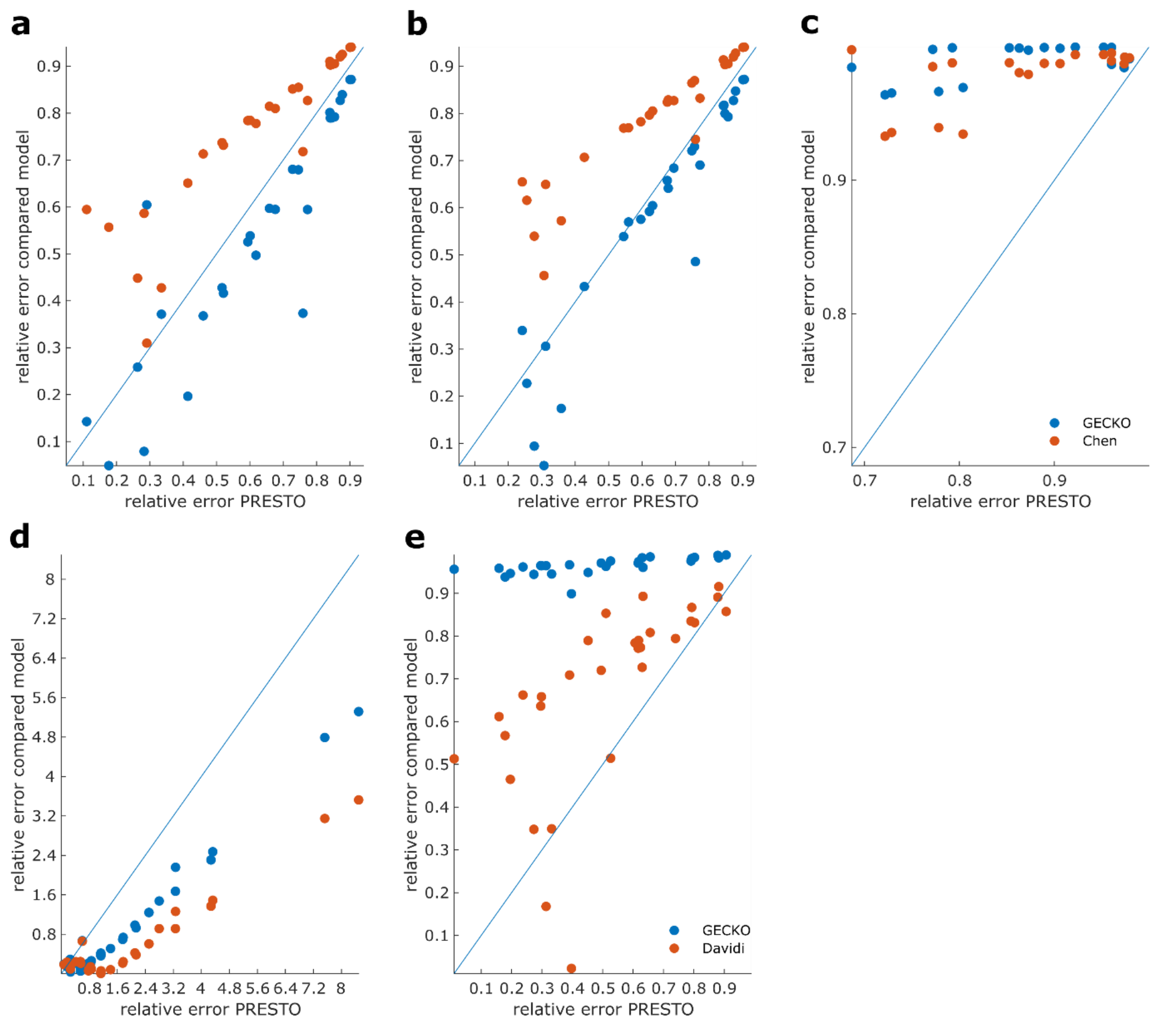


**Fig S10. Comparison of predicted growth from $k_{cat}$ corrected based on PRESTO, maximum correction from GECKO, and estimated from pFBA.** $k_{cat}$ estimates obtained from pFBA studies of *S. cerevisiae* (Chen & Nielsen, 2021) (**a-c**) and *E. coli* (Davidi *et al*, 2016) (**d,e**) were used to correct initial BRENDA values in the raw pcGEM. To generate a condition-independent GECKO model the maximum $k_{cat}$ over all conditions was used. The y-axis denotes relative error of the GECKO (blue) or pFBA (red) model in comparison to the relative error of the PRESTO model, plotted on the x-axis. **(a,d)** Only the measured total protein pool was used to constrain the solution and condition-specific uptake rates were bounded by 1000 $\frac{mmol}{h\ gDW}$; **(b)** in *S. cerevisiae* available measured uptake rates were also considered **(c,e)** in addition to the previous constraints abundances of enzymes measured in all conditions were used as constraints. The compared pcGEMs in each condition used the same respective biomass coefficients, GAM, $\sigma$, and $P_{tot}$ values (Methods).

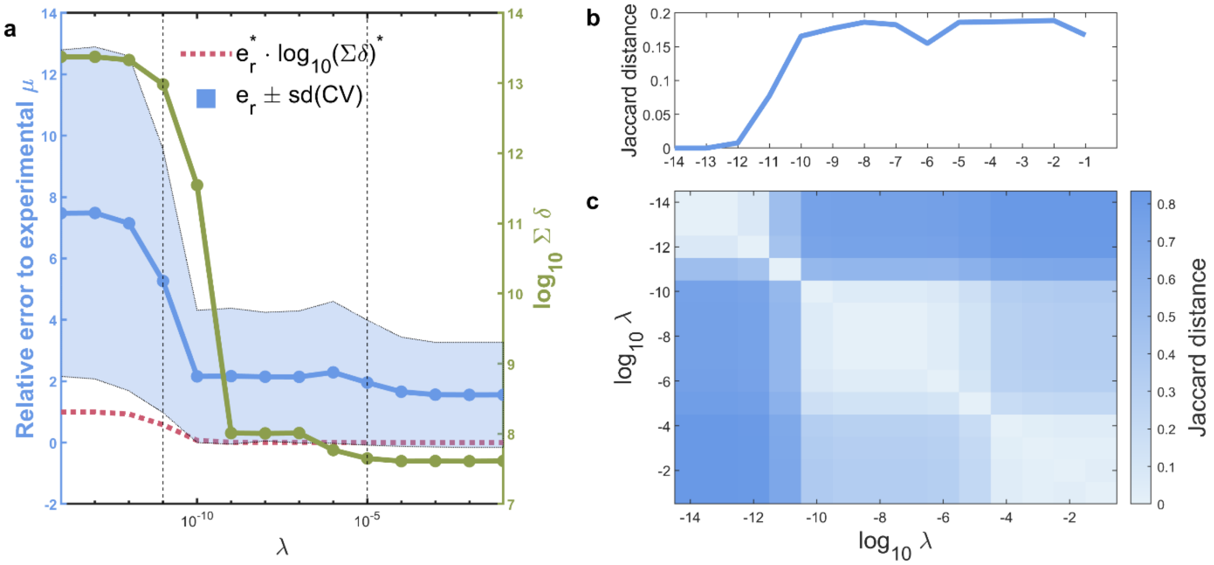


**Fig S11. Cross validation to obtain the optimal value for parameter λ for *E. coli* pcGEM. (a)** Shown are the average relative error ($e_r$, blue solid line), average sum of added corrections δ (green solid line), and the scoring metric that was used to find the optimal value (red dotted line). Altogether, 14 values for λ were explored in the range from $10^{-14}$ to $10^{-1}$. The optimal value for λ was determined by the first inflection point of the scoring metric (right dashed vertical line). The left dashed vertical line represents the value at which the sum of δ reaches a plateau. **(b)** Average Jaccard distance between cross-validation folds over ten iterations for each of the explored λ. **(c)** Average Jaccard distance between the union of corrected $k_{cat}$ values over ten iterations for each pair of explored λ values.

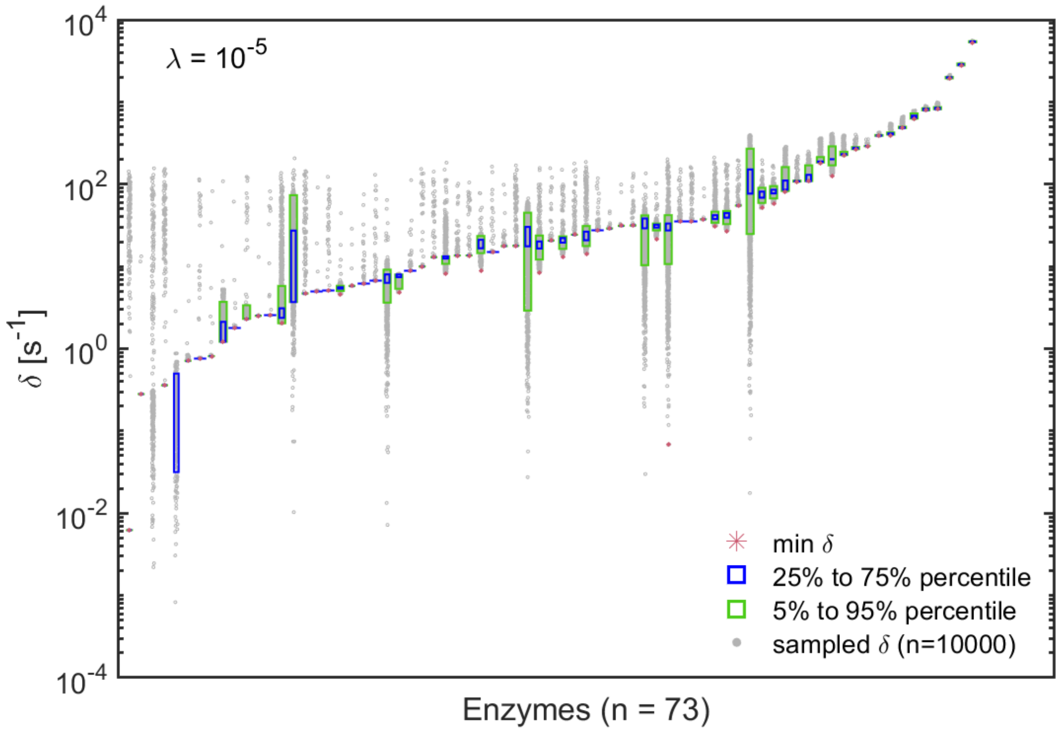


**Fig S12. Precision of $k_{cat}$ corrections introduced by PRESTO using the optimal λ value for the *E. coli* pcGEM.** The minimum and maximum values for each δ were determined by variability analysis in which the relative errors and the sum of corrections, δ, are fixed to values obtained from PRESTO. Using these ranges, 10,000 random points were sampled uniformly across the feasible region.

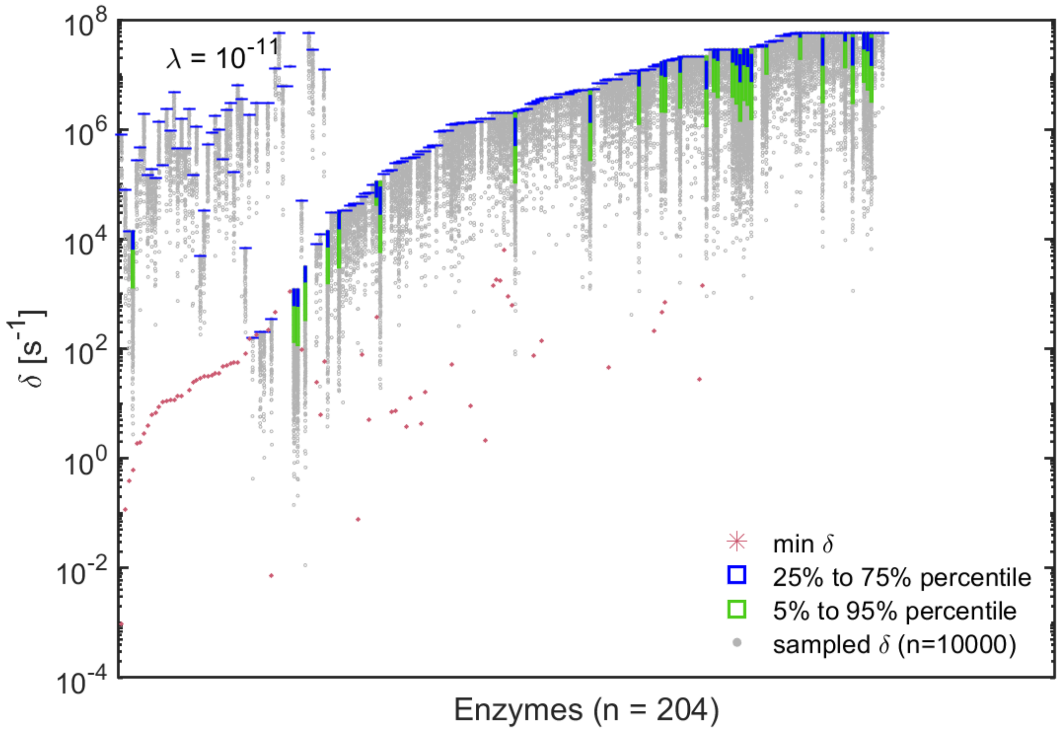


**Fig S13. Precision of $k_{cat}$ corrections introduced by PRESTO using the λ value for which the average sum of corrections plateaus for the *E. coli* pcGEM.** The minimum and maximum values for each δ were determined by variability analysis in which the relative errors and the sum of corrections, δ, are fixed to values obtained from PRESTO. Using these ranges, 10,000 random points were sampled uniformly across the feasible region. To arrive at feasible optimization programs, the feasibility tolerance for the solver was decreased to $10^{-6}$).

**a**

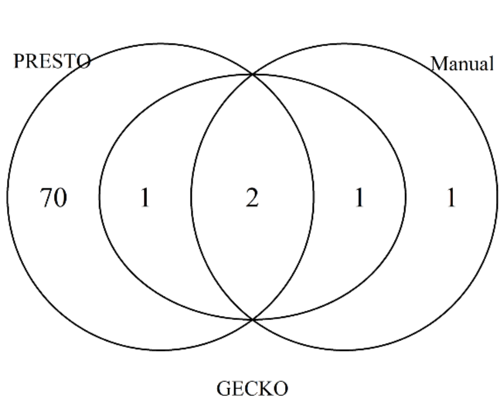


**b**

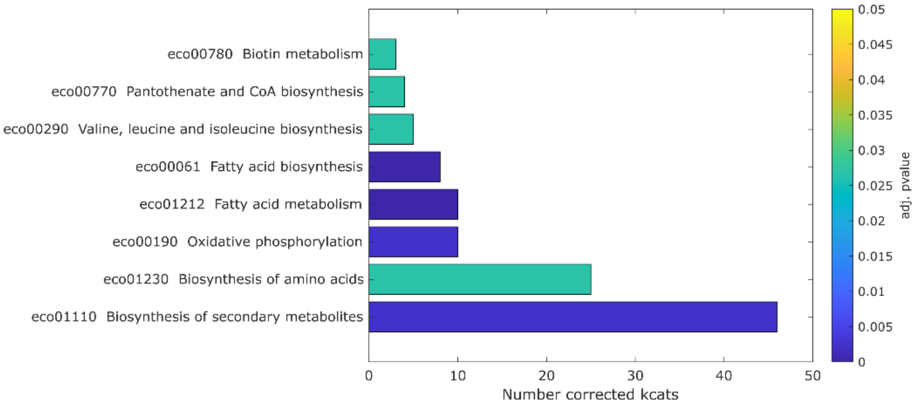


**Fig S14. Enzymes with $k_{cat}$ values corrected by PRESTO in *E. coli*. (a)** Venn diagram showing the overlap of enzymes whose $k_{cat}$ values were manually corrected (Sánchez *et al*, 2017) (“Manual”), automatically corrected by the GECKO heuristic in any of the conditions (“GECKO”) or corrected by PRESTO (“PRESTO”, $\lambda = 10^{-5}$) **(b)** KEGG Pathway terms significantly enriched in the set of enzymes corrected by PRESTO in the *E.coli* model. The x-axis gives the number of enzymes with corrected $k_{cat}$ values linked to the given term. The one-sided p-value was calculated using the hypergeometric density distribution and was corrected using the Benjamini-Hochberg procedure (Benjamini & Hochberg, 1995).

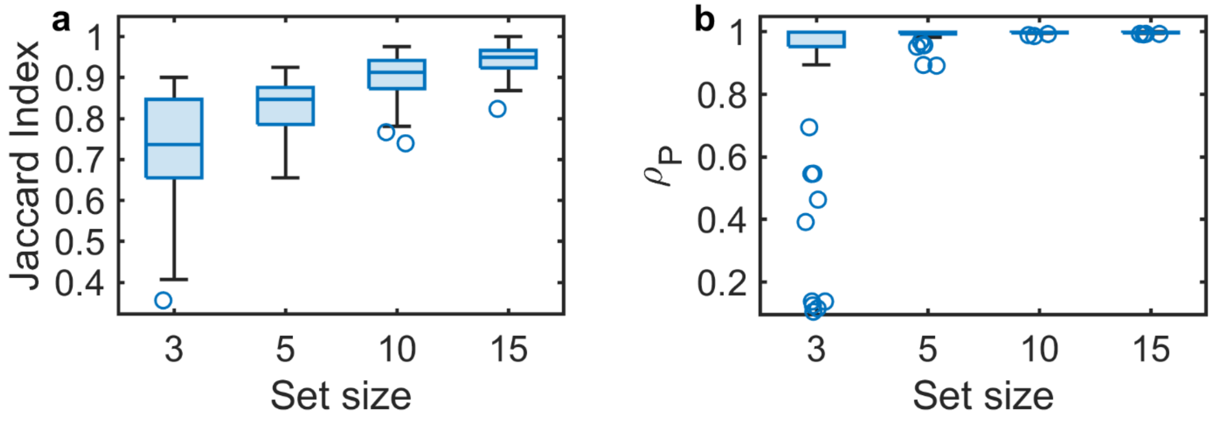


**Fig S15. Robustness analysis of turnover number corrections for *S. cerevisiae*.** The PRESTO approach as used to determine turnover number corrections with different sizes of randomly selected subsets (n=50) of the full dataset comprising 27 conditions. The sets of enzymes with corrected $k_{cat}$ values were compared to the solution obtained with all conditions. This was done by **(a)** Jaccard Index of the protein identifiers and **(b)** Pearson correlation of the corrections $\delta$ for enzymes whose turnover numbers were corrected using the subset and the full set of conditions.

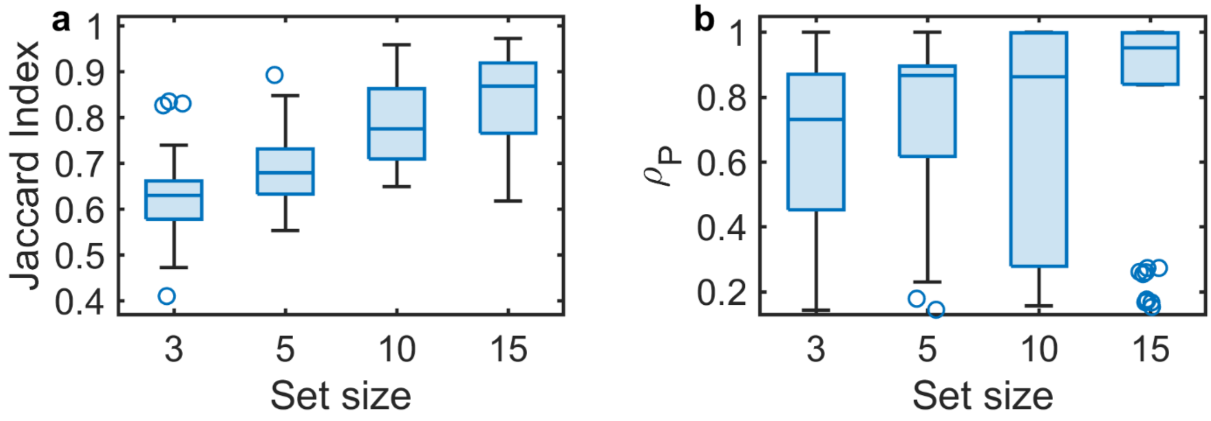


**Figure S16. Robustness analysis of turnover number corrections for *E. coli*.** The PRESTO approach as used to determine turnover number corrections with different sizes of randomly selected subsets (n=50) of the full dataset comprising 27 conditions. The sets of enzymes with corrected $k_{cat}$ values were compared to the solution obtained with all conditions. This was done by **(a)** Jaccard Index of the protein identifiers and **(b)** Pearson correlation of the corrections $\delta$ for enzymes whose turnover numbers were corrected using the subset and the full set of conditions.

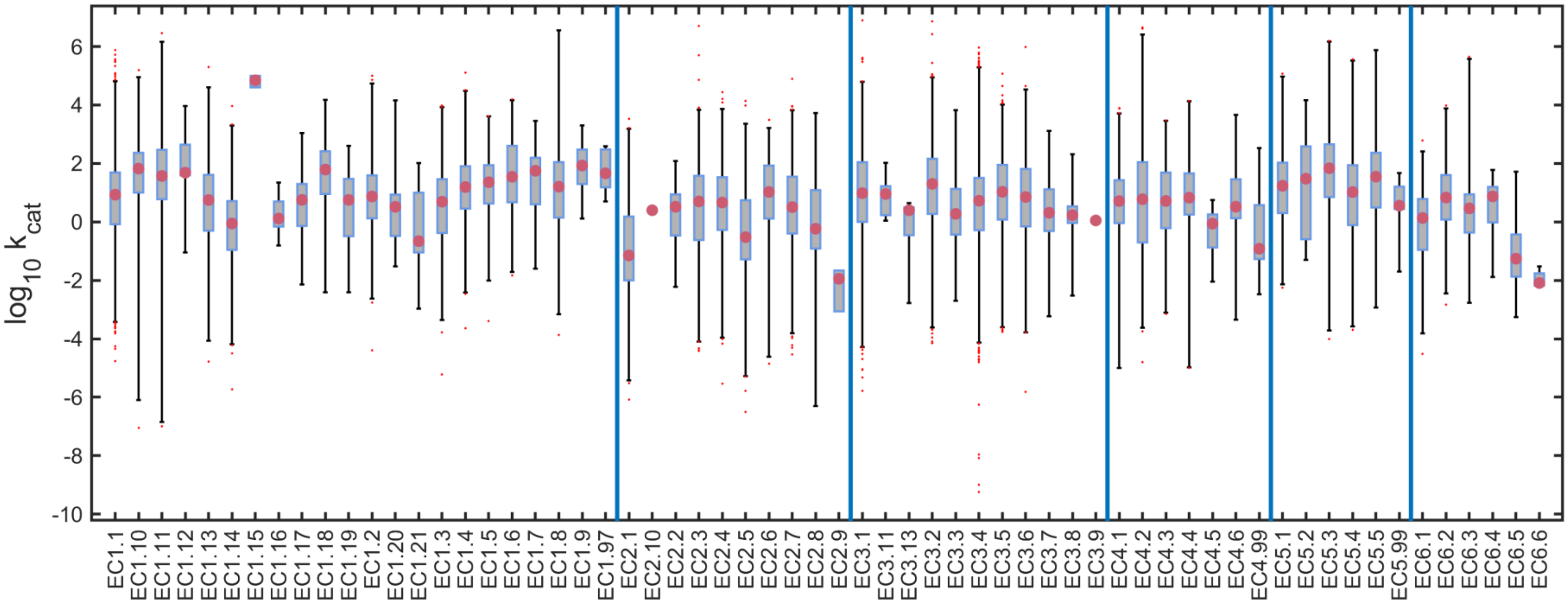


**Fig S17. Distribution of turnover numbers within EC number classes in BRENDA.** The boxplots show $k_{cat}$ values for six major EC classes and respective sub-classes as contained in the $k_{cat}$ file provided with the GECKO toolbox (Sánchez *et al*, 2017). Red dots depict median values, boxes indicate interquartile ranges and whiskers cover 99.3% of data points. Outliers are shown as red stars. Blue vertical lines provide separation between major EC classes.

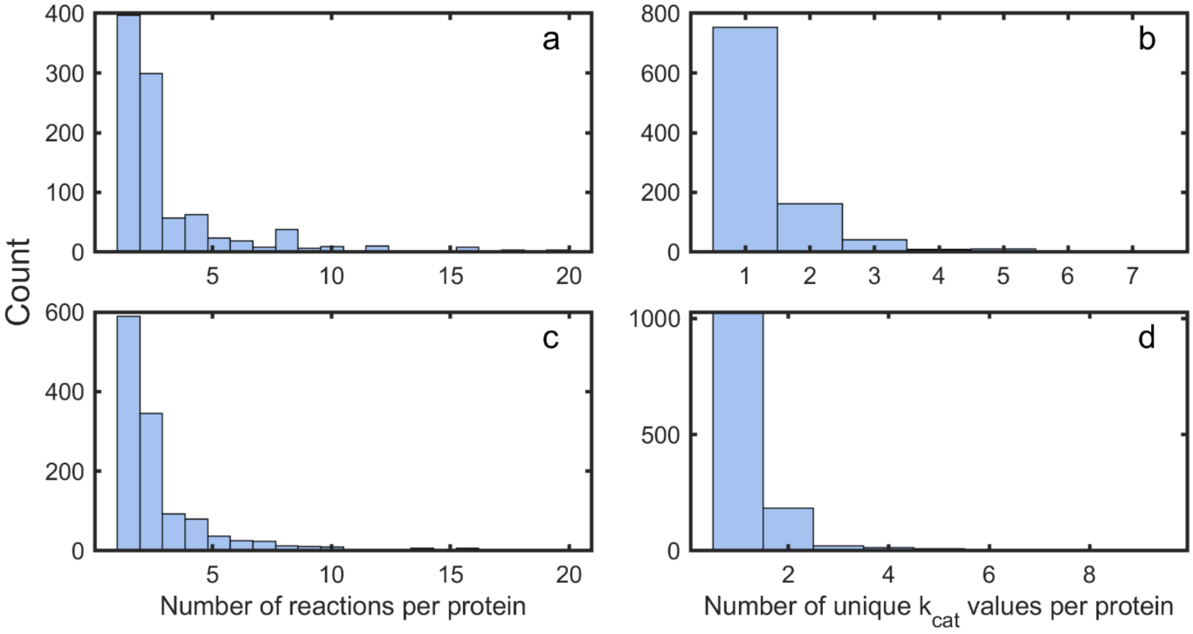


**Fig S16. Numbers of reactions per enzyme and number of unique $\mathbf{k_{cat}}$ values per enzyme in the *S. cerevisiae* and *E. coli* pcGEMs. (a,c)** Histograms of the number of reactions per enzyme (i.e., promiscuity). The histograms were plotted with a range between 1 and 20 reactions per protein. For *S. cerevisiae*, this threshold was exceeded by 31 proteins with a maximum of 384 reactions (*E. coli*: 14 above threshold, maximum: 42) **(b,d)** Histograms showing the number of unique $k_{cat}$ values per enzyme.

**Legend for Table S1 (separate file)**

**Table S1. Model performance and $k_{cat}$ corrections. A** Model performance only using experimental uptake rates and total protein abundance but no protein pool as constraints in *S. cerevisiae* pcGEM. Relative prediction error of growth rate by PRESTO model and average relative prediction error and standard deviation of GECKO models. NaN indicates models that were infeasible. **B** Values of corrected turnover numbers from PRESTO ('PRESTO_KCAT') and considering the maximum of all $k_{cat}$ corrections from condition-specific GECKO models ('GECKO_KCAT') in *S. cerevisiae*. **C** Values of corrected turnover numbers from PRESTO ('PRESTO_KCAT') and considering the maximum of all $k_{cat}$ corrections from condition-specific GECKO models ('GECKO_KCAT') in *E. Coli*. **D** Information from uniprot und KEGG servers for the enzymes whose turnover numbers were corrected by GECKO and PRESTO in S. cerevisiae (see also Fig3b and FigS16). **E** Information from uniprot und KEGG servers for the enzymes whose turnover numbers were corrected by both GECKO and PRESTO in E. coli (see also FigS12a).

**Table S2. Minimal medium used for simulations with the eciML1515 model for *E. coli*.**

| Symbol | Exchange reaction |
|---|---|
| $Na^+$ | EX_na1_e_REV |
| $P_i$ | EX_pi_e_REV |
| $Cl^-$ | EX_cl_e_REV |
| $K^+$ | EX_k_e_REV |
| $NH4^+$ | EX_nh4_e_REV |
| $Mg^{2+}$ | EX_mg2_e_REV |
| $SO_4^{2-}$ | EX_so4_e_REV |
| $MoO_4^{2-}$ | EX_mobd_e_REV |
| $Mn^{2+}$ | EX_mn2_e_REV |
| $Ni^{2+}$ | EX_ni2_e_REV |
| $Zn^{2+}$ | EX_zn2_e_REV |
| $Cu^{2+}$ | EX_cu2_e_REV |
| $Ca^{2+}$ | EX_ca2_e_REV |
| $Fe^{2+}$ | EX_fe2_e_REV |
| $Fe^{3+}$ | EX_fe3_e_REV |
| $Cd^{2+}$ | EX_cd2_e_REV |
| $Co^{2+}$ | EX_cobalt2_e_REV |
| $H_2O$ | EX_h2o_e_REV |
| $O_2$ | EX_o2_e_REV |
